# Social Determinants modulate NK cell activity via obesity, LDL, and DUSP1 signaling

**DOI:** 10.1101/2023.09.12.556825

**Authors:** Yvonne Baumer, Komudi Singh, Andrew S. Baez, Christian A. Gutierrez-Huerta, Long Chen, Muna Igboko, Briana S. Turner, Josette A. Yeboah, Robert N. Reger, Lola R. Ortiz-Whittingham, Christopher K.E. Bleck, Valerie M. Mitchell, Billy S. Collins, Mehdi Pirooznia, Pradeep K. Dagur, David S.J. Allan, Daniella Muallem-Schwartz, Richard W. Childs, Tiffany M. Powell-Wiley

## Abstract

Adverse social determinants of health (aSDoH) are associated with obesity and related comorbidities like diabetes, cardiovascular disease, and cancer. Obesity is also associated with natural killer cell (NK) dysregulation, suggesting a potential mechanistic link. Therefore, we measured NK phenotypes and function in a cohort of African-American (AA) women from resource-limited neighborhoods. Obesity was associated with reduced NK cytotoxicity and a shift towards a regulatory phenotype. *In vitro*, LDL promoted NK dysfunction, implicating hyperlipidemia as a mediator of obesity-related immune dysregulation. Dual specific phosphatase 1 (DUSP1) was induced by LDL and was upregulated in NK cells from subjects with obesity, implicating DUSP1 in obesity-mediated NK dysfunction. *In vitro*, DUSP1 repressed LAMP1/CD107a, depleting NK cells of functional lysosomes to prevent degranulation and cytokine secretion. Together, these data provide novel mechanistic links between aSDoH, obesity, and immune dysregulation that could be leveraged to improve outcomes in marginalized populations.

## Introduction

Obesity, defined by the World Health Organization as excessive or abnormal accumulation of adipose tissue, is associated with distinct health risks and has a complex pathogenesis involving biological, environmental, socioeconomic, and psychosocial factors.^1^ Social determinants of health (SDoH) play a crucial role in the development of obesity; these SDoH include neighborhood environment, socioeconomic status (SES), education, and healthy food access.^2–4^ Furthermore, adverse SDoH (aSDoH) can impede the treatment of individuals with obesity and magnify the racial and ethnic disparities seen in obesity and related comorbidities.^5^ After age-adjustment, obesity is most prevalent in non-Hispanic Black populations followed by Hispanic and non-Hispanic White populations, with non-Hispanic Black women at highest risk for incident obesity.^6,7^

Obesity increases the risk of cardiovascular disease (CVD), hyperlipidemia, type 2 diabetes, and some cancers ^8–15^, reducing life expectancy by 5-20 years.^8^ Many of these outcomes have been linked to excess obesity-related inflammation. Increased visceral adiposity promotes the release of pro-inflammatory mediators including tumor necrosis factor α (TNFα) and interleukin (IL)-6.^16, 17^ These adipokines, cytokines, and chemokines modulate hematopoiesis^18^ and immunity^13, 19–23^ to promote adverse obesity-related and aSDoH-related outcomes.

Obesity can affect the phenotype and function of almost any immune cell subset.^24^ Many prior studies have focused on monocytes ^25, 26^ and neutrophils ^27, 28^ due to their critical role in atherogenesis.^29–31^ Yet natural killer (NK) cells – a subset of innate-like lymphocytes with cytolytic function – may also regulate atherogenesis^32, 33^. NK cells are also important for early host-pathogen responses, cancer surveillance, immunomodulation, and cellular cytotoxicity.^34^ NK functional subsets are categorized by CD56 expression, with highly proliferative CD56^bright^CD16^-^ NK cells acting as immunomodulators, while CD56^dim^CD16^+^ NK cells are more cytotoxic.^35^

Class III obesity (body mass index [BMI] ≥40 kg/m^2^) has been associated with metabolic reprogramming of NK cells, reduced numbers of cytotoxic CD3^-^/CD56^+^ NK cells, and reduced NK cell function.^36^ In children with obesity and insulin resistance, reduced NK cell numbers were also accompanied by defective tumor lysis.^37^ Adipose tissue-resident NK cells can express lower levels of activating NKp30 and NKp44 receptors in individuals with obesity.^38^ Together, these findings suggest that obesity modulates NK cell distribution and function.^36, 39–41^ Yet the specific pathways modulating NK cell alterations in individuals with obesity are incompletely characterized. This is especially true in marginalized populations experiencing aSDoH, who are traditionally underrepresented in research studies.

Herein, we examined the impact of obesity on the phenotype and function of circulating NK cells. To specifically investigate the effects of aSDoH-related obesity, we studied NK cells in a cohort of African American (AA) women from resource-limited neighborhoods, comparing individuals with vs. without overweight/obesity. As we have previously demonstrated that low density lipoprotein (LDL) regulates immune cell function, we selected pathways modulated by both LDL and obesity for further analysis. We hypothesized that these pathways might be central to obesity-mediated changes in NK phenotype and function, promoting CVD, cancer, and other obesity-related comorbidities.

## Results

### The frequency of NK cell subsets in blood of AA females as related to obesity and aSDoH

Between AA females from resource-limited neighborhoods who did (O/O, BMI ≥ 25 kg/m^2^) or did not (NO/O, BMI < 25 kg/m^2^) have overweight/obesity (**Table S1**),), there was no significant difference in the frequency of overall lymphocytes (44.7 ± 4.8% versus 40.1 ± 5.1%), CD19^+^ B cells (4.9 ± 0.8% versus 3.1 ± 0.5%), CD3^+^/CD56^-^ T cells (31.9 ± 3.5% versus 30.4 ± 4.6%), CD3^+^ expressing CD56 T cells (1.9 ± 0.7% versus 1.3 ± 0.4%), and CD3^-^/CD56^+^ NK cells (4.2 ± 0.5% versus 4.2 ± 0.7%) within circulating CD45^+^ mononuclear cells **(Fig. 1A)**. We observed a significantly higher proportion of CD56^bright^/CD16^dim^ NK cells (4.6 ± 0.8% versus 2.7 ± 0.4%; p=0.029) and significantly lower proportion of CD56^dim^/CD16^+^ NK cells (88.5 ± 1.5% versus 92.1 ± 1.0%; p=0.048) within the overall CD3^-^/CD56^+^ NK cell population in O/O AA females compared to age-matched NO/O AA females (**Fig. 1B).**

**Figure 1:**
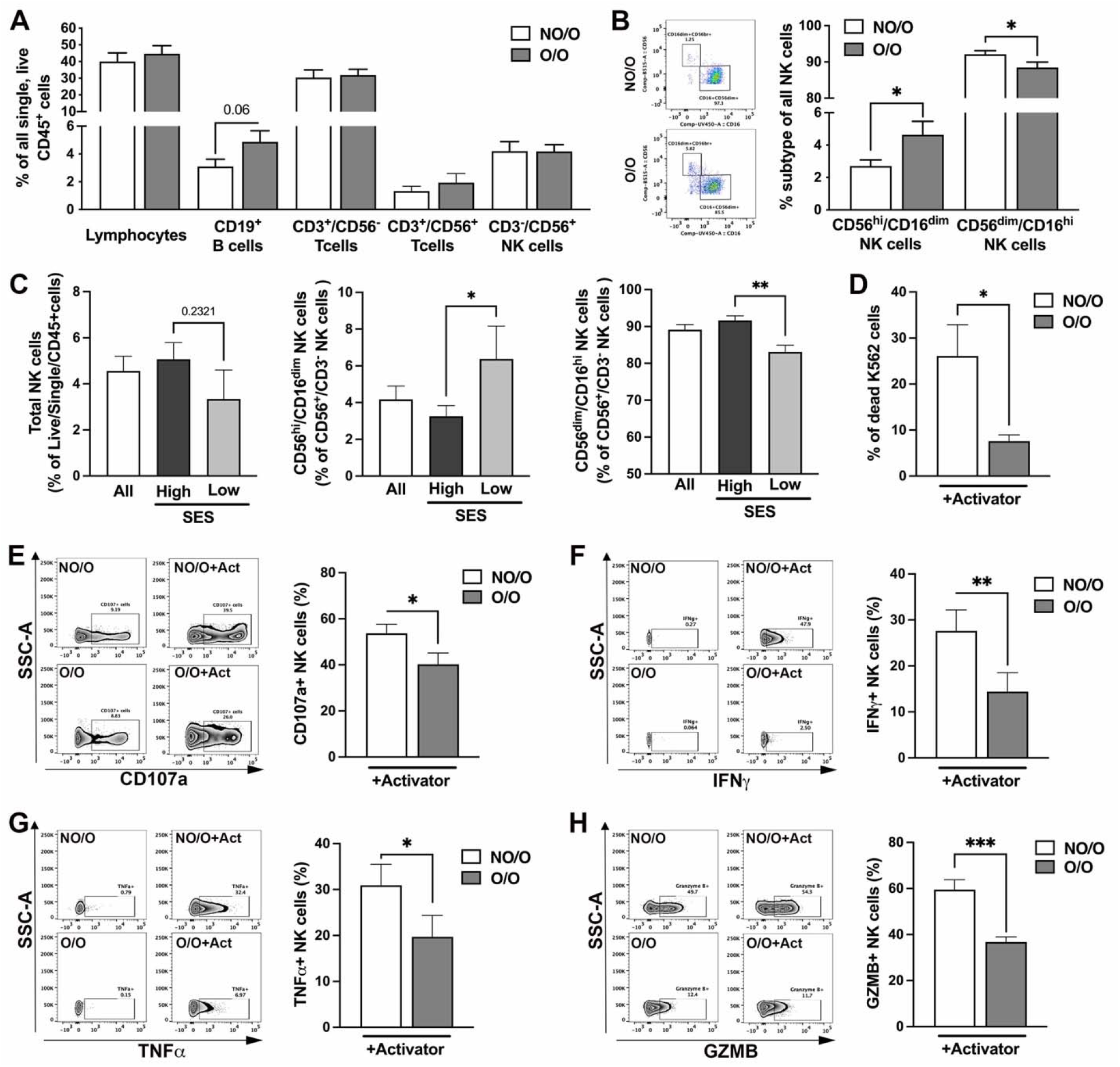
Obesity and lower socioeconomic status associated with differences in NK cell subset profiles. Study participants were AA females from resource-limited neighborhoods who did (O/O; BMI>25, n=14) or did not present with overweight/obesity (NO/O; BMI<25, n=15), respectively. (**A/B**) Whole blood was examined by flow cytometry to analyze the lymphocytes, namely, CD19^+^B cells, CD3^+^/CD56^-^ T cells, CD3^+^/CD56^+^ T cells, and CD3^-^/CD56^+^ NK cell populations. **(B)** Deeper phenotyping of CD56+ NK cells by CD16 expression allows a general identification of CD56^hi^/CD16^dim/-^ and CD56^dim^/CD16^hi^ NK cell phenotypes. (Mann-Whitney test comparing two patient groups for each immune cell population) **(C)** Lower SES is defined as annual household income <$60,000 based on median household income in Washington, DC. (**C**) Total NK cells and each NK cell subset by SES. (**D-F**) NK cells were isolated from AA females who did (O/O; BMI>25, n=13) or did not present with overweight/obesity (NO/O; BMI<25, n=15) and subsequently were subjected to the degranulation assay. By using flow cytometry, the proportion of dead K562 cells (**D**) as a measure of NK cell killing ability under activating conditions, the expression of CD107a (**E**), as a measure of NK cell degranulation, and the intracellular expression of IFNγ (**F**), TNFα (**G**), and granzyme B (**H**) were detected. (**D/F/G**: Mann-Whitney test was performed to determine statistical significance; **D/E/H**: Unpaired T-test was performed to determine statistical significance); (Significance was established when a p-value < 0.05 was present comparing individual groups with the data appropriate test; *indicates significance between groups; Flow data are accompanied by representative dot or volcano blots.)

Within this cohort of AA females, we also examined associations between SDoH (i.e. neighborhood social environment factors, individual-level SES, psychosocial factors) and total, CD56^bright^/CD16^dim/-^, and CD56^dim^/CD16^+^ NK cells (**Table 1**, **Fig. 1C**). Among neighborhood social environment factors, neither neighborhood deprivation index (NDI) as a marker of neighborhood socioeconomic deprivation, perceived neighborhood physical/social environment, nor perceived neighborhood social cohesion were associated with detectable differences in NK cell distribution. However, higher individual-level SES as measured by household income associated with higher CD56^dim^/CD16^+^ NK cell proportion in both the unadjusted and adjusted models (**Table 1**, **Fig. 1C**). Among the psychosocial factors, higher social isolation was associated with lower NK total populations independent of BMI (β=-0.63, p=0.02). Higher social isolation also associated with lower CD56^dim^/CD16^+^ NK cell proportions (β=-0.77, p=0.01; β=-0.69, p=0.02 in BMI or BMI/ASCVD adjusted models, respectively). Higher depressive symptoms trended towards a significant association with lower total NK cells in the BMI and the fully adjusted models (β=-0.46, p=0.06; β=-0.47, p=0.09; in BMI or BMI/ASCVD adjusted models, respectively). This association was likely driven by the CD56^dim^/CD16^+^ NK cell proportions (β=-0.22, p=0.04; β=-0.54, p=0.05; in BMI or BMI/ASCVD adjusted models, respectively). Within the depressive symptom sub-types, higher sadness-related depressive symptoms were associated with lower CD56dim/CD16+ NK cell proportions and higher sleep-related depressive symptoms were associated with lower total NK cell proportions (**Table 1**).

**Table 1A:** Multivariable Linear Regression Modeling of Associations Between Neighborhood Social Environment, Individual-Level Socioeconomic Status, and Psychosocial Factors as Chronic Stressors and NK Cell Populations in Cohort of AA Females^†^. Models are unadjusted (Model 1), adjusted for BMI (Model 2), and adjusted for ASCVD 10-yr risk score and BMI (Model 3). Findings displayed as standardized beta coefficients (p-values).

|  | Total NK Cells |  |  | CD56<sup>dim</sup>/CD16<sup>+</sup> NK Cells |  |  | CD56<sup>bright</sup>/CD16<sup>dim/-</sup> NK Cells |  |  |
| --- | --- | --- | --- | --- | --- | --- | --- | --- | --- |
|  | Model 1 | Model 2 | Model 3 | Model 1 | Model 2 | Model 3 | Model 1 | Model 2 | Model 3 |
| **NDI<sup>^</sup>** | 0.17<br>(0.51) | 0.27<br>(0.33) | 0.16<br>(0.61) | 0.04<br>(0.87) | 0.16<br>(0.59) | 0.07<br>(0.83) | 0.17<br>(0.52) | 0.06<br>(0.86) | 0.19<br>(0.51) |
| **Neighborhood Physical /Social Environment<sup>#</sup>** | 0.09<br>(0.76) | 0.16<br>(0.58) | 0.37<br>(0.30) | -0.31<br>(0.26) | -0.23<br>(0.52) | 0.21<br>(0.57) | 0.27<br>(0.33) | 0.15<br>(0.68) | 0.12<br>(0.77) |
| **Neighborhood Social Cohesion<sup>&</sup>** | 0.47<br>(0.08) | 0.46<br>(0.09) | 0.54<br>(0.10) | -0.03<br>(0.91) | -0.06<br>(0.84) | -0.08<br>(0.81) | 0.04<br>(0.88) | 0.10<br>(0.77) | 0.14<br>(0.71) |
| **SES<sup>‡</sup>** | 0.42<br>(0.09) | 0.50<br>(0.06) | 0.50<br>(0.11) | **0.51**<br>**(0.04)\*** | **0.67**<br>**(0.01)\*** | **0.65**<br>**(0.02)\*** | -0.29<br>(0.27) | -0.52<br>(0.10) | -0.47<br>(0.16) |
| **Social Isolation** | **-0.59**<br>**(0.03)\*** | **-0.63**<br>**(0.02)\*** | -0.54<br>(0.08) | **-0.54**<br>**(0.04)\*** | **-0.77**<br>**(0.01)\*** | **-0.69**<br>**(0.02)\*** | 0.32<br>(0.27) | 0.56<br>(0.11) | 0.45<br>(0.21) |
| **Depressive Symptoms and Symptom Sub-types:** |  |  |  |  |  |  |  |  |  |
| **Overall Depressive Symptoms<sup>§</sup>** | -0.35<br>(0.20) | -0.46<br>(0.06) | -0.47<br>(0.09) | -0.22<br>(0.43) | **-0.22**<br>**(0.04)\*** | **-0.54**<br>**(0.05)\*** | -0.17<br>(0.54) | 0.15<br>(0.63) | 0.11<br>(0.76) |
| **Sadness<sup>§</sup>** | -0.40<br>(0.14) | **-0.50**<br>**(0.049)\*** | -0.49<br>(0.09) | -0.32<br>(0.24) | **-0.65**<br>**(0.01)\*** | **-0.62**<br>**(0.02)\*** | -0.08<br>(0.79) | 0.25<br>(0.45) | 0.18<br>(0.59) |
| **Anhedonia/Loss of Interest<sup>§</sup>** | -0.48<br>(0.07) | -0.53<br>(0.06) | -0.50<br>(0.11) | -0.28<br>(0.31) | -0.42<br>(0.18) | -0.38<br>(0.24) | -0.04<br>(0.89) | 0.05<br>(0.89) | -0.05<br>(0.90) |
| **Lack of Appetite<sup>§</sup>** | 0.43<br>(0.11) | 0.36<br>(0.16) | 0.40<br>(0.19) | **0.52**<br>**(0.049)\*** | 0.40<br>(0.16) | 0.39<br>(0.19) | -0.34<br>(0.22) | -0.14<br>(0.66) | -0.11<br>(0.74) |
| **Lack of Sleep<sup>§</sup>** | **-0.66**<br>**(0.007)\*\*** | **-0.74**<br>**(0.001)\*\*** | **-0.84**<br>**(0.001)\*\*** | -0.29<br>(0.28) | -0.54<br>(0.06) | -0.53<br>(0.09) | -0.05<br>(0.86) | 0.18<br>(0.60) | 0.14<br>(0.70) |
| **Change in Thinking<sup>§</sup>** | -0.19<br>(0.48) | -0.26<br>(0.36) | -0.40<br>(0.25) | -0.05<br>(0.86) | -0.18<br>(0.57) | -0.20<br>(0.55) | -0.21<br>(0.43) | -0.16<br>(0.64) | -0.16<br>(0.68) |
| **Guilt<sup>§</sup>** | -0.11<br>(0.69) | -0.15<br>(0.62) | -0.09<br>(0.79) | -0.33<br>(0.10) | -0.31<br>(0.29) | -0.49<br>(0.13) | 0.01<br>(0.98) | 0.15<br>(0.67) | 0.10<br>(0.79) |
| **Tired/Fatigue<sup>§</sup>** | -0.01<br>(0.96) | -0.12<br>(0.65) | -0.03<br>(0.93) | -0.08<br>(0.77) | -0.40<br>(0.14) | -0.37<br>(0.18) | -0.22<br>(0.43) | 0.11<br>(0.71) | 0.04<br>(0.88) |
| **Movement/Agitation<sup>§</sup>** | 0.33<br>(0.22) | 0.23<br>(0.33) | 0.16<br>(0.40) | 0.28<br>(0.32) | 0.02<br>(0.95) | -0.02<br>(0.93) | -0.25<br>(0.37) | 0.12<br>(0.67) | 0.21<br>(0.49) |

<sup>†</sup> N=17 of 29 AA women (NO/O vs. O/O) for whom psychosocial and environmental factors were available.
<sup>^</sup> NDI - neighborhood deprivation index derived from U.S. Census data (higher score=more socioeconomically deprived neighborhood)
<sup>&</sup> Perceptions about neighborhood physical/social environment measured by validated survey (higher score=greater perceived resources in the physical/social environment)
<sup>#</sup> Perceived neighborhood social cohesion measured by validated survey (higher score=more neighborhood cohesion)
<sup>‡</sup> SES – individual-level socioeconomic status (higher value=higher self-reported household income)
<sup>||</sup> Social Isolation - measured by validated survey (higher score=increasing social isolation)
<sup>§</sup> Depression and depression sub-scores were measured by validated surveys (higher score=more depression, sadness, lack of appetite, lack of sleep, increased perception of guilt)

### Obesity is associated with reduced NK cell degranulation and cytokine secretion

Because obesity was associated with an increase in regulatory CD56^bright^/CD16^dim/-^ cells, we compared cytotoxicity of NK cells from O/O and NO/O subjects. NK cells isolated from AA females with O/O displayed a 71% decreased ability to kill target K562 cells compared to NK cells from AA females with or without O/O (**Fig. 1D**). Additionally, surface expression of CD107a, a marker of degranulation, was 25% lower on NK cells of AA females with O/O as compared to females without O/O (**Fig. 1E**). Under NK cell activating conditions and after contact with susceptible K562 target cells, we detected significantly lower levels of intracellular granzyme B (34% decrease), IFNγ (52% decrease), and TNFα (64% decrease) in NK cells from AA females with O/O compared to NO/O AA females (**Fig. 1F**). We did not see significant changes in intracellular perforin or GM-CSF production (S**uppFig. 1C/D**).

### Impact of LDL on NK cell function

To understand how obesity might impact NK cell function; we utilized an *ex vivo* approach. We examined the influence of serum from a cohort of AA individuals (n=60) on healthy donor NK cells (Cohort 2, **Table S2**). After 24 hours of treatment the degranulation assay towards K562 cells was performed. Subsequently these results were subjected to multivariable regression analysis with the levels of various serum components in these samples (**Table 2, Table S3, Fig. 2A/B**). No significant associations were found between levels of HDL or ApoA1 and NK cell degranulation as measured by CD107a expression. Higher LDL and ApoB levels associated with lower CD107a expression among study participants on no lipid-lowering therapy (**Table 2**, **Fig. 2A/B**). We also found that higher serum total cholesterol, LDL, and ApoB were associated with lower intracellular IFNγ expression independent of ASCVD risk, BMI, and SES among participants on no lipid-lowering therapy.

**Figure 2:**
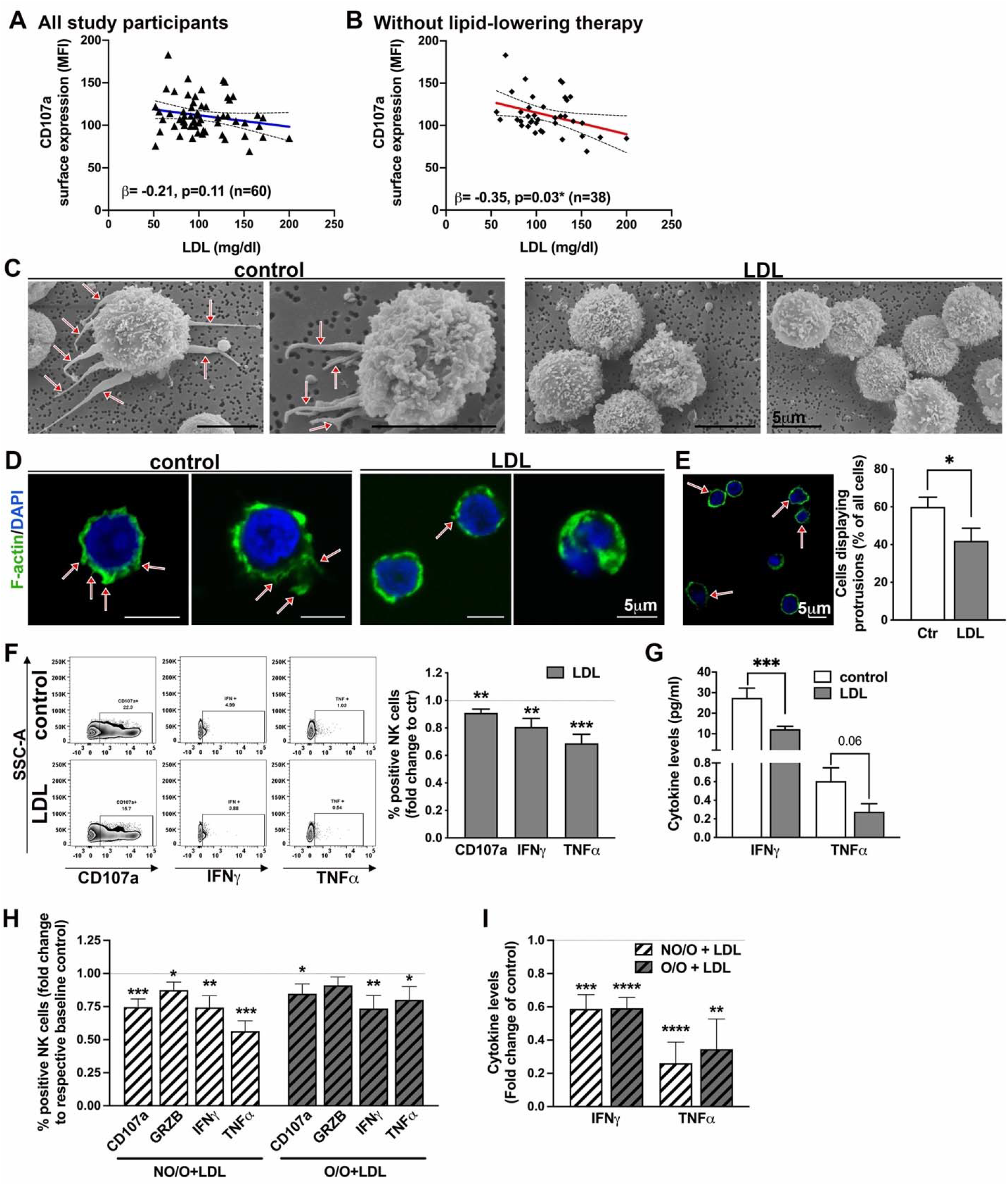
Obesity is associated with NK cell dysfunction potentially driven by LDL. **(A/B)** NK cells were isolated from a healthy blood bank donor, treated over night with study participants’ serum (n=60), and subsequently subjected to the degranulation assay towards K562 cells. Surface CD107a expression levels were utilized in multivariable regression analysis against LDL levels of the study participants. (A: Data from all 60 study participants, B: display of regression curve from participants without lipid-lowering therapy (subgroup n=38)). (**C-G**) Freshly isolated primary NK cells from healthy bloodbank donors were treated over night with LDL (50mg/dl) or vehicle control. **(C)** Utilizing scanning electron microscopy, the presence of protrusions (arrows) was examined as a sign of NK cell activity (n=3). (**D/E**) Labeling the actin cytoskeleton of vehicle or LDL treated NK cells using FITC-Phalloidin with subsequent confocal microscopy imaging allowed for quantification of cells presenting protrusions. Graph in (G) displays the result of n=9. Statistical significance was tested using the unpaired T-test. (**F/G**) Degranulation assay towards K562 cells was performed after overnight treatment with subsequent flow cytometry analysis of surface CD107a, and intracellular cytokine expression (F, n=22). Supernatants were subjected to ELISA-based measurement of secreted cytokines (G) (n=17). (**H/I**) NK cells were isolated from AA females with (O/O) or without (NO/O) overweight/obesity (BMI>25, n=13 and BMI<25, n=15), treated with LDL overnight and subsequently subjected to the degranulation assay towards K562 cells (**H**) and the levels of secreted cytokines determined by ELISA (**I**). (Significance was established when a p-value < 0.05 was present comparing individual groups with an unpaired t-Test for each vehicle/LDL dataset; *indicates significance between groups; F is accompanied by a representative volcano blots.)

**Table 2:**
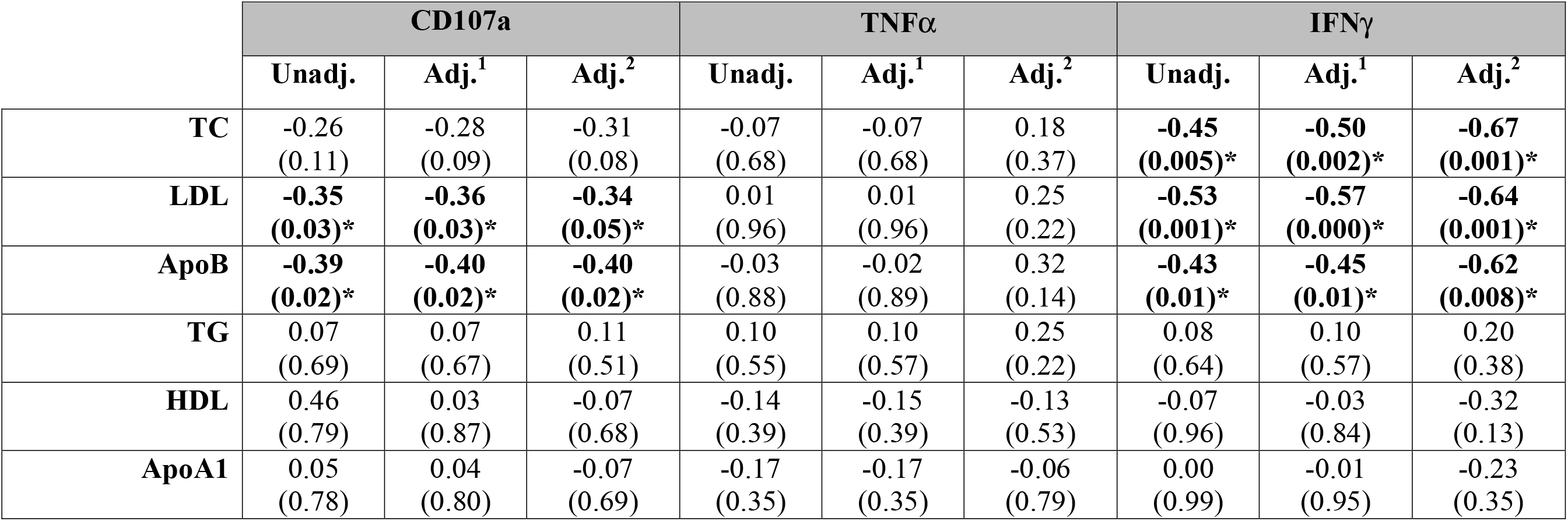
In an *ex vivo* approach healthy donor primary NK cells were incubated with donor serum (patients summarized in Supplementary Table 2) for 24hours and afterwards subjected to the degranulation assay. Regression associations between detected CD107a surfaces expression as well as intracellular TNFα and IFNγ expression and clinical parameters in unadjusted, fully adjusted model 1 (adjusted for ASCVD and BMI) and adjusted model 2 (adjusted for ASCVD, BMI, SES) are shown as standardized ß-value with the corresponding p-value in parenthesis in individuals without lipid-lowering therapy.

Subsequently, we examined a potential effect of LDL (500mg/ml = 50mg/dl) on primary freshly isolated NK cells from blood bank donors and subsequently analyzed phenotypic and functional changes (**Fig. 2C-G**). First, we utilized scanning electron microscopy (SEM) to examine NK cell morphology and protrusion presence, as these are crucial for the cytotoxic capabilities of NK cells.^42^ We found that LDL-treated NK cells displayed fewer protrusions compared to vehicle-treated NK cells (**Fig. 2C**). These findings were confirmed utilizing z-stack confocal microscopy of F-actin labeled NK cells (**Fig. 2D/E**). Next, we measured NK cell degranulation and cytokine production following LDL exposure (**Fig. 2F/G**). LDL exposure decreased CD107a surface expression (p=0.001) compared to vehicle treatment. Intracellular NK cell IFNγ and TNFα were significantly decreased by LDL exposure (p=0.003 and p=<0.0001 respectively). Secretion of IFNγ was significantly decreased by LDL exposure (p=0.0006), with a similar but non-significant trend in TNFα secretion (p=0.06) (**Fig. 2G**).

To determine if these effects of LDL on healthy volunteer NK cells are also occurring in NK cells isolated from our study participants of AA females with NO/O or O/O (Cohort 1), we exposed study participants NK cells with LDL and performed the degranulation assay (**Fig. 2H/I**). Both groups of NK cells displayed decreased surface CD107a expression and decreased intracellular expression of IFNγ and TNFα (**Fig. 2H**). LDL also significantly decreased levels of secreted IFNγ and TNFα in the supernatant of NO/O and O/O NK cells (**Fig. 2I**).

### Identification of DUSP1 involvement in NK cell dysfunction related to obesity and LDL

To discover pathways connecting obesity and LDL to NK cell function, we took an unbiased approach. We compared the transcriptomic changes induced by LDL in healthy volunteer NK cells to the differentially expressed genes (DEG) in NO/O versus O/O (Cohort 1) subjects (**Fig. 3A**). Principal component analyses (PCA) NO/O versus O/O cells, or control versus *in vitro* LDL-treated cells revealed clustering of the samples by type (Fig. **S2A/B**). 1645 DEG were identified in NO/O versus O/O, with 1126 upregulated and 519 downregulated (**Fig. S2C)**. A custom select subset of pathways of interest was applied for pathway enrichment analysis. Enrichment analysis for immunometabolic pathways revealed significant enrichment for cytokine signaling, IFNγ-mediated signaling, fatty acid metabolism and lysosome processes (**Fig. 3B**). 8621 DEG were seen in control versus LDL-treated NK cells, with 2498 upregulated and 6122 downregulated (**Fig. S2D**). Enrichment analysis for immunometabolic pathways showed significant enrichment in cytokine and lipid metabolism (**Fig. 3C**).

**Figure 3:**
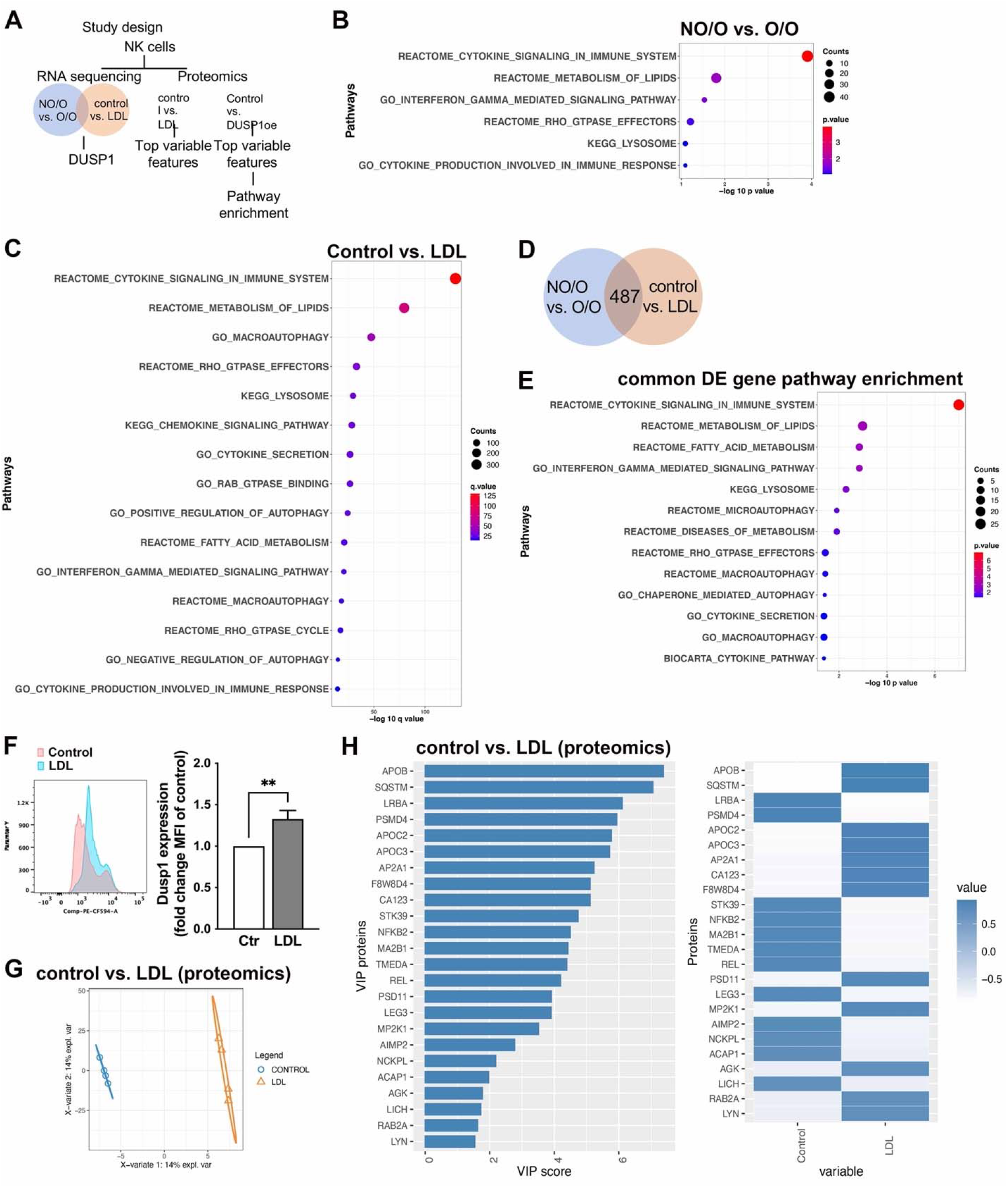
Comparative analysis of two RNA sequencing sets reveals *DUSP1* as an overlapping significantly upregulated gene, known to negatively regulate inflammatory processes. **(A)** A schema of the unbiased omics analysis undertaken in this study. The NK cells from the indicated human sub-groups or after LDL exposure were used for gene expression profiling or for proteomics as indicated. Subsequent downstream analysis helped identify common targets and led to the identification of Dusp1. Freshly isolated primary NK cells from 1) either AA women with or without O/O (dataset 1 1: n=5 each group) or 2) healthy donors treated with vehicle or LDL overnight (dataset 2: n=4) were subjected to RNAsequencing analysis. Vehicle/LDL treated NK cells were additionally subjected to proteomics analysis (n=4). (**B-C**) Top 15 pathways enriched by differentially expressed genes in either NO/O vs O/O (B) or control vs. LDL (C) NK cells presented as a dot plot. Each dot is a pathway labeled on y-axis, the size of the dot is scaled to the number of DE genes overlapping with the pathway and are plotted on negative log 10 q value on the x-axis. **(D)** Venn diagram showing overlap of differentially expressed genes identified in the NO/O vs. O/O comparison and control vs. LDL comparison. 487 genes were commonly differentially expressed genes between both datasets. **(E)** Top 15 pathways enriched by 487 commonly differentially expressed genes shown in D is presented as a dot plot. Each dot is a pathway labeled on y-axis, the size of the dot is scaled to the number of DE genes overlapping with the pathway and are plotted on negative log 10 q value scale on the x-axis. **(F)** Flow cytometry analysis of Dusp1 protein expression in vehicle or LDL exposed healthy donor NK cells. (n=6, unpaired t-Test) **(G)** Partial least square discriminant analysis (PLSDA) plot of the proteomics data from the control and LDL exposed samples. **(H)** Variables important in predicting the control and LDL exposed NK cells. The VIP score of the indicated proteins on the y axis and their mean expression in control and LDL treated NK cells on the x axis is presented.

To identify genes commonly perturbed in NK cells from AA females with O/O and in *in vitro* LDL-exposed NK cells, DE genes from both datasets were analyzed together (**Fig. 3D**). A total of 487 genes were significantly differentially expressed in both datasets. Of these genes, 132 were concordantly upregulated and 148 were concordantly repressed in both datasets. Pathway enrichment analysis of these concordantly DE genes showed enrichment of cytokine as well as IFNγ-mediated signaling, fatty acid metabolism, lysosome, and autophagy processes (**Fig. 3E, Fig. S2E/F**). Notably, the gene encoding dual specificity protein phosphatase 1 (*DUSP1*) was highly induced by both obesity and LDL (NO/O versus O/O: logFC=2.8 p=0.00009 and control versus LDL: logFC=0.2 p=0.008). Interestingly, NK cell *DUSP1* gene expression significantly correlated with BMI (R^2^=0.73, p=0.02) (**Fig. S2G/H**) and correlation with LDL-c (concentration) and LDL-p (particle number) (p=0.06) trended towards significance in study participants not on lipid-lowering therapy.

DUSP1 (MKP-1) is the first characterized dual-specific phosphatase able to de-phosphorylate and inactivate all three MAPKs (JNK, p38, ERK). As an inducible DUSP-family member, it is a negative feedback regulator of NFκB signaling. Loss of DUSP1 leads to increased expression of LPS-induced TNFα, IL-6, or IFNγ of various cell types, highlighting its role in repression of inflammation. Furthermore, Dusp1 regulates proliferation, autophagy, metabolic homeostasis, and glucocorticoid signaling. ^43^

To examine if LDL-exposure of healthy volunteer NK cells not only affected *DUSP1* mRNA expression but also its protein expression, we performed flow cytometry analysis of intracellular Dusp1 expression. LDL-exposure of healthy volunteer NK cells resulted in 32.9±10.1% increased expression of DUSP1 protein (p=0.009) (**Fig. 3F)**.

Our finding that LDL and obesity highly induced *DUSP1* in NK cells led us to hypothesize that DUSP1 might mediate lipid/obesity-induced NK functional changes. To test this hypothesis, we analyzed public NK cell transcriptomic datasets for associations between *DUSP1* expression and NK cell function. *DUSP1* expression decreased by a mean of 18.7% (p=0.27) in NK cells activated with IL-12 (GSE50838)^44^, a strong inducer of NK cell cytotoxicity. Additionally, *DUSP1* was 2-fold downregulated (p<0.0001) after NK cell expansion using the Amplification and Expansion System (NKAES) with irradiated K562mbIL21 feeder cells in the presence of IL-2 (GSE128696)^45^ (**Fig. S2I/J**).

Proteomic analysis (LC-MS/MS) of LDL-exposed NK cells identified 45 differentially induced and 34 differentially repressed proteins. NFKB2, a protein critically involved in DUSP1-regulation of NFκB signaling^46^, was significantly decreased, consistent with LDL-mediated *DUSP1* induction. (**Fig. 3G/H**). Additional variables of importance in prediction (VIP) in LDL-exposed cells as included autophagy protein p62 (SQSTM1), and proteasomal complex proteins PSMD4 and PSD11 (**Fig. 3H**).

To assess the role of DUSP1 in NK cell function more directly, we utilized two experimental approaches. First, we overexpressed DUSP1 in the NK92 NK cell line. Overexpression decreased expression of CD107a by 30.4±3% and intracellular TNFα by 25.3±7% (**Fig. 4A/B**), as well as inhibiting IFNγ and TNFα release. We then assessed the impact of pharmacologic DUSP1 inhibition on LDL-mediated NK cell functional changes (**Fig. 4C-E**). DUSP1 inhibition rescued LDL-repression of CD107a, intracellular TNFα expression, and TNFα secretion (**Fig. 4C,D**). Scanning electron microscopy revealed that DUSP1 inhibition also rescued LDL-driven changes in NK cell morphology (**Fig. 4E**).

**Figure 4:**
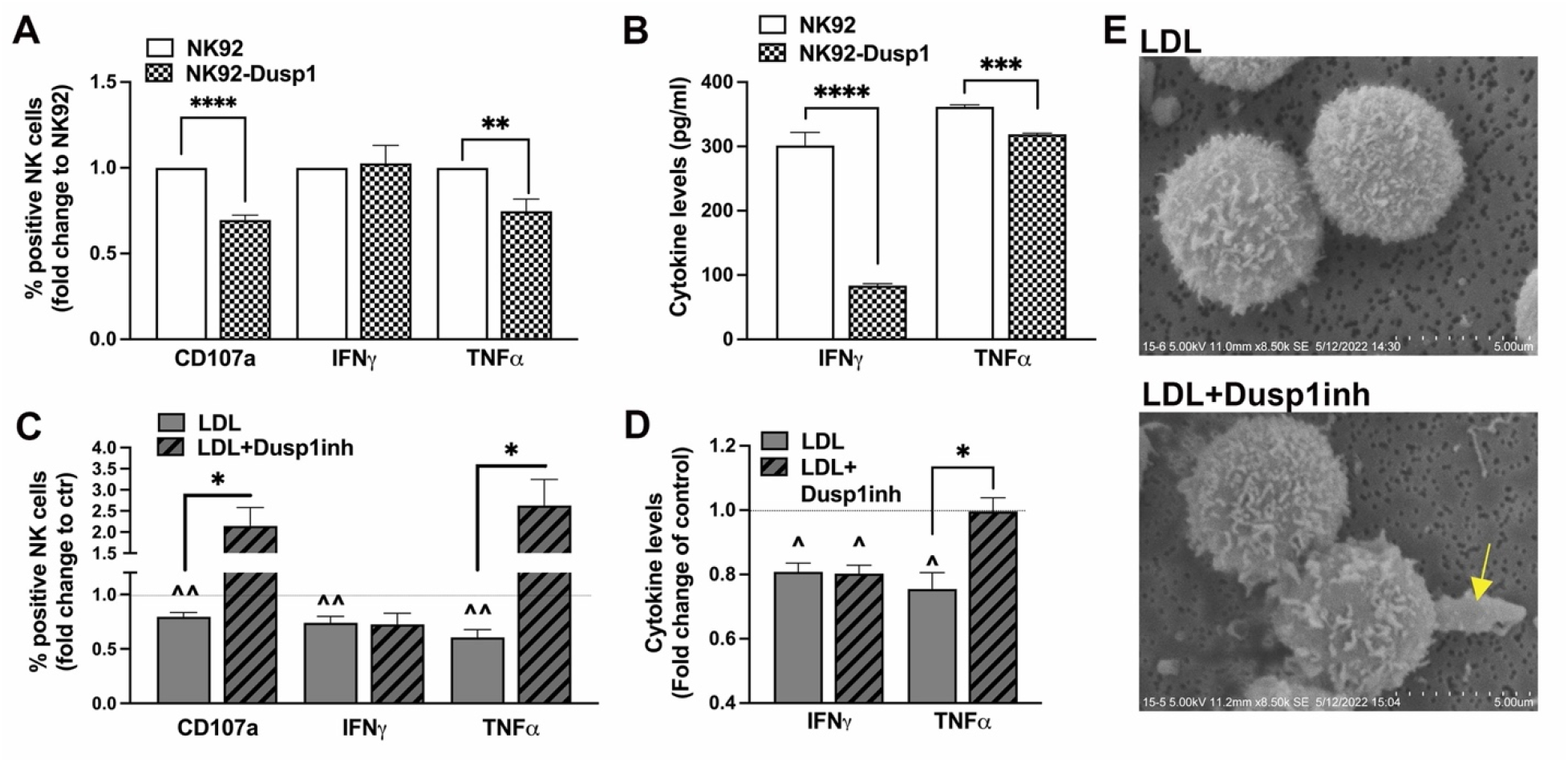
Dusp1 is an important regulator in NK cell function. (**A/B**) Dusp1 was overexpressed in NK92 cells. Experiments are carried out comparing empty vector (EV) control NK92 cells to Dusp1 overexpressing NK92 cells. After subjecting the two cell lines to K562 cells flow cytometry analysis was performed to determine degranulation (CD107a) and intracellular cytokine presence. Cytokine secretion was measured using ELISA as displayed in (B). (n=6 each, unpaired t-Test for all, except Mann-Whitney for TNFα dataset) (**C-E**) Freshly isolated primary NK cells are treated with LDL in absence or presence of a Dusp1 inhibitor and subjected to the degranulation assay (C) with cytokine release analysis (D). (n=9, RM-One Way Anova with Tukey correction) **(E)** SEM imaging of LDL and LDL+Dusp1 inhibitor treated NK cells (n=3). (Significance was established when a p-value < 0.05 was present comparing individual groups; *indicates significance between groups; ^indicates significance to vehicle control treatment)

### Downstream targets of DUSP1 in NK cells from individuals with obesity or LDL-exposed NK cells

DUSP1 is a known negative feedback regulator of cytokine production via its impact on MAPK and NFκB activity.^46^ To determine if DUSP1 regulates these pathways in NK cells, we determined the impact of DUSP1 inhibition on NFκB pathway-related protein expression in LDL-exposed healthy volunteer NK cells (**Fig. S3**). LDL exposure reduced NFκB protein visualized by immunofluorescence staining; this effect was abrogated by the DUSP1 inhibitor (**Fig. S3A**). Interestingly, LDL exposure did not significantly alter pathway-related protein activity, but reduced NFκB activity by 51±1% when compared to vehicle control (**SuppFig. 3B**). This LDL-induced reduction in NFκB activity was not observed in the presence of the DUSP1 inhibitor (fold change to control: LDL 0.50±0.1 versus LDL+Dusp1inh 1.39±0.4).

### DUSP1 promotes lysosomal dysfunction in NK cells

To further explore the impact of *DUSP1* on NK cells, an unbiased gene expression and proteomics profiling of the DUSP1 overexpressing NK92 cell line compared to the parental line was performed (**Fig. 5A-D, Fig. S4**). DE and PCA analyses showed that samples clustered based on the DUSP1 overexpression status (**Fig. S4A**). A large number of upregulated and downregulated genes were observed (**Fig. S4B**). Pathway enrichment analysis of the upregulated and downregulated DE genes suggested perturbations in lysosome, autophagy, and cytokine secretion processes (**Fig. 5A/B**). To validate these findings and examine the impact of DUSP1 overexpression on lysosomes, RT-qPCR and flow cytometry were performed. DUSP1 overexpressing NK cells display 59% reduced *LAMP1* mRNA levels, while *TFEB*, a transcription factor previously reported to regulate LAMP1 (CD107a) expression^47^, was not affected (**Fig. 5C**). There was a 47% reduction in intracellular LAMP1 (CD107a) protein by flow cytometry (**Fig. 5D**), accompanied by reduced lysosomal vesicles on TEM (**Fig. 5E**).

**Figure 5:**
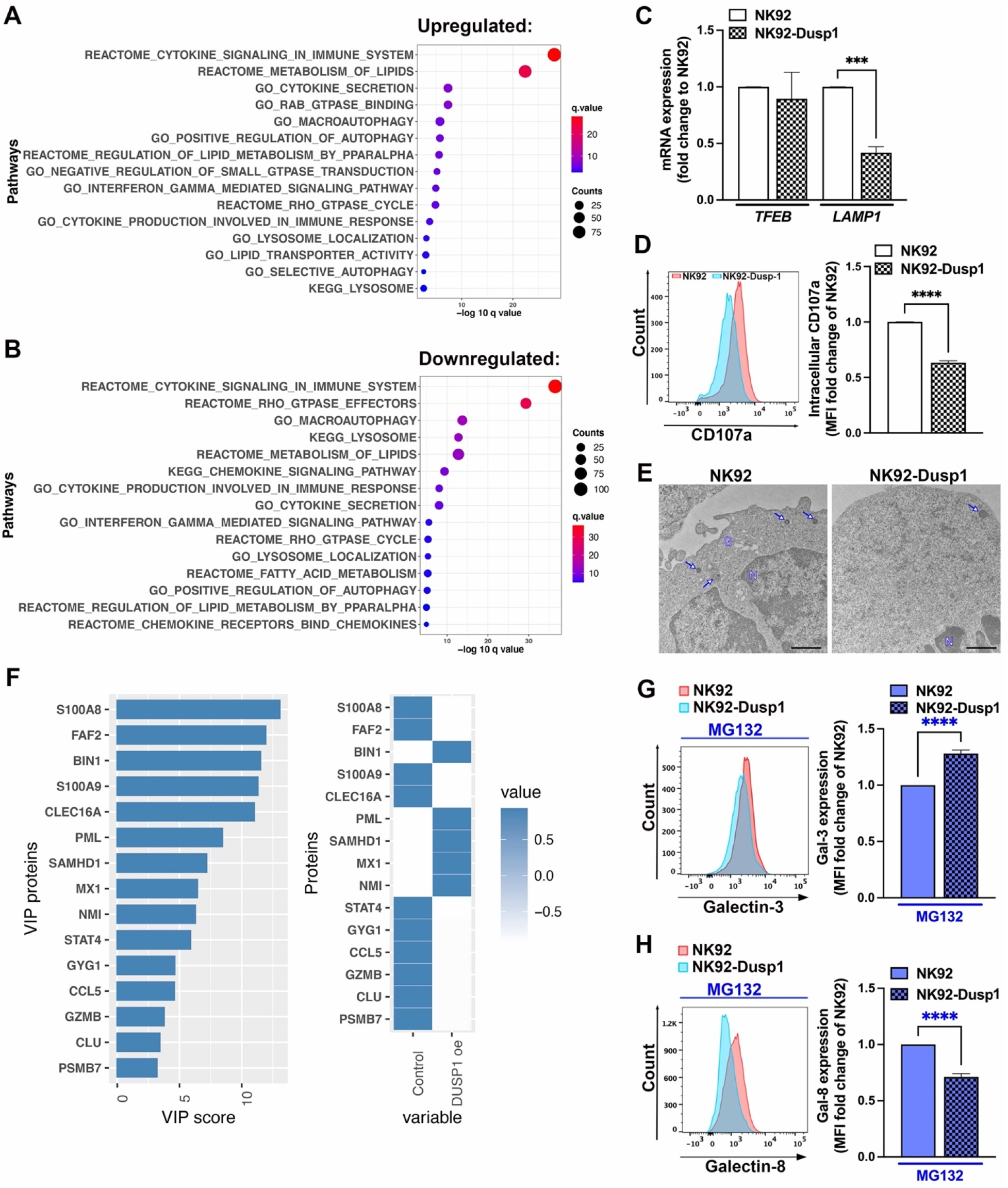
Dusp1 overexpression impacts lysosome biology. Dusp1 was overexpressed in NK92 cells, a well-accepted NK cell line. The created cell line and its control were examined to better understand pathways regulated by Dusp1. **(A/B)** Top 15 pathways enriched by the differentially expressed genes that were either upregulated (A) or downregulated (B) in Dusp1 oe cells is presented as dot plot. Each dot is a pathway labeled on y-axis, the size of the dot is scaled to the number of DE genes overlapping with the pathway and are plotted on negative log 10 q value scale on the x-axis. (**C-E**) Dusp1 overexpressing NK cells and their control were examined for intracellular Lamp-1 expression using flow cytometry (C, n=6, unpaired t-Test), *LAMP1* and *TFEB* mRNA expression (n=3, unpaired t-Test), and TEM imaging (n=3 pooled, scale bar 1μm). **(E)** Variables important in predicting the control and Dusp1 overexpressing cells. The VIP score of the indicated proteins on the y axis and their mean expression in control and Dusp1 oe cells on the x axis is presented. (**G/H**) Galectin 3/8 expression were analyzed in both cell lines using flow cytometry in the presence of the proteasome inhibitor MG132. (n=10, G-Mann Whitney test, H-unpaired t-Test) (Significance was established when a p-value < 0.05 was present comparing individual groups; *indicates significance between groups)

To verify if these findings also apply for LDL-treated primary NK cells, a set of similar experiments was performed in presence or absence of the DUSP1 inhibitor (**Fig. S4 C-G**). Treatment with LDL did not significantly affect *LAMP1* mRNA but addition of the DUSP1 inhibitor increased TFEB as well as *LAMP1* mRNA compared to LDL exposure alone (**Fig. S4C**). Additionally, we found that LAMP1 protein expression was decreased after LDL exposure, which could be prevented by concurrent addition of the DUSP1 inhibitor (**Fig. S4D/E**). Since lysosomal location and positioning within a cell impacts their pH^48^ and is crucial for cytolytic function of cytotoxic T lymphocytes^49^ and cytokine secretion is dependent on a variety of lysosomal vesicles^50^, we quantified the number of lysosomes and examined vesicle location by TEM (**Fig. S4F/G**). LDL exposure reduced the number and altered the location of lysosomal vesicles of various morphologies^51^, supporting intracellular trafficking impairment and potentially explaining reduced cytokine secretion from NK cells with LDL exposure (**Fig. S4F/G**). Inhibition of DUSP1 ameliorated the impact of LDL as more lysosomes were found closer to the cell periphery.

To better understand the LDL-induced, DUSP1-dependent loss of NK cell lysosomes, we performed proteomic analysis of the DUSP1 overexpressing NK cell line (**Fig. 5F, Fig. S4H**). S100A8, shown to mediate inflammation in NK cells, and GZMB, important for NK cell cytotoxicity, were downregulated in DUSP1-overexpressing cells. These observations are consistent with the dysregulation of cytokine secretion processes identified via gene expression profiling. Pathway analysis (**Fig. S4I**) supported the findings on TEM, showing impact on pathways regulating secretory granules, degranulation, and lysosomal transport. Pathways involved in proteasome-dependent lysophagy were also suggested to be dysregulated.

Prior studies have shown a relationship between various lipids, including LDL, and lysosomal damage^52, 53^; lysosomal damage has been associated with lysosomal repair processes, lysophagy, or replenishment in a galactin3/8-dependent manner.^54^ We examined whether DUSP1 overexpression might affect lysosome numbers by changing galectin-3/8 expression and affecting lysosome homeostasis. We found that galectin-3 expression is increased in DUSP1 overexpressing cells by 28.1±3.1% while galectin-8 expression is decreased by 28.8±2.9%, suggesting a switch to lysophagy without lysosome replenishment (**Fig. 5G/H**).

## Discussion

The associations between aSDoH and obesity are well-documented and represent a major source of disparities in both CVD and cancer morbidity and mortality. However, the underlying mechanisms and subsequent consequences on immune cell function and intracellular signaling are incompletely understood. In this study, we focused on the potential effects of aSDoH and obesity on NK cells in a high-risk population of AA women disproportionately impacted by aSDoH. In this population, lower SES, higher social isolation and greater depressive symptoms were associated with a pathological shift in NK cell immunophenotype. NK cells of AA women with obesity also displayed reduced degranulation, cytolysis, and cytokine secretion. In individuals not on lipid-lowering therapy, higher LDL was associated with NK cell dysfunction, suggesting a mechanistic link to obesity. Accordingly, LDL promoted a dysfunctional NK cell phenotype that recapitulated the immunophenotype observed in NK cells from individuals with obesity. Unbiased omics-based analyses identified DUSP1 as a crucial regulator of obesity- and LDL-mediated NK cell dysfunction. This was further supported by the effects of DUSP1 inhibition and overexpression on basal NK cell function and LDL-induced NK cell dysfunction. Together, these results strongly suggest that in individuals exposed to aSDoH, obesity promotes NK cell dysfunction through LDL and DUSP1.

Prior studies have examined the relationship between aSDoH and NK cells, but these studies have often been done in animal models^55, 56^, or potentially confound results by not providing results from minoritized populations or individuals at increased risk for CVD.^57, 58^ In rhesus macaques, changes in social status and harassment significantly altered NK and T-helper immunophenotypes, thereby regulating responses to viral infection.^56^ aSDoH can also promote sex-specific differences in immune function; for example, higher childhood SES may be associated with increased NK cell cytotoxicity in young men without obesity (mean age 20 years with no chronic medical issues and BMI<30 kg/m^2^), but not in young women without obesity.^59^ By contrast, we have described reduced proportions of circulating CD56^dim^/CD16^hi^ cytotoxic NK cells in healthy African American blood bank donors compared to healthy Caucasian blood bank donors.^60^ Additionally, and in contrast to these prior studies^59^, our results highlight the potential importance of psychosocial stress and socioeconomic conditions on NK cell function in women from under-resourced neighborhoods. These findings are relevant to middle-aged women with CVD risk factors, a population at particularly high risk for adverse CVD outcomes.^61^

Our findings suggesting a relationship between psychosocial stress and NK cell dysfunction are consistent with prior studies associating psychosocial factors, including chronic stress, anxiety^62^, and depression^63^, with NK cell function. For example, stress and anxiety were associated with reduced CD56^+^/CD16^+^ NK cells in a cohort of patients with chronic tinnitus.^62^ NK cytotoxicity, degranulation, and IFN-γ secretion were also reduced in individuals who had experienced maternal deprivation as a form of early life adversity. ^64^ Maternal deprivation was associated with increased serum glucose levels, a marker of cardiometabolic dysfunction and a risk factor for CVD.^64^ Of note, in our cohort of middle-aged adult women with cardiometabolic risk factors, NK cell dysfunction was associated socioeconomic level and psychosocial stress even after multivariable adjustments for BMI and ASCVD 10-year risk score. Future studies will be needed to explore the relationships between psychological stress, cardiometabolic diseases like pre-diabetes and diabetes, and NK cell dysfunction in aSDoH-affected populations.

To our knowledge, this is the first study to specifically investigate obesity-related NK changes in AA women experiencing chronic aSDoH. From epidemiologic studies, aSDoH, including neighborhood deprivation^65^, neighborhood crime and disorder^66^, lower socioeconomic resources^67^, depression^68^, and perceived stress^69^, are associated with incident obesity, especially among AA women who are disproportionally impacted by aSDoH and subsequent obesity.^70^ An extensive body of prior work has shown that obesity negatively affects NK cell phenotype and function.^39,71^ Similarly, we observed a switch in NK cell immunophenotype towards a CD56^bright^/CD16^dim/-^ population, which is associated with reduced cytotoxicity. Accordingly, we detected reduced degranulation, surface CD107a expression, and IFNγ production.^36^ More work is needed to determine if the NK cell immunophenotypic changes observed in this study may occur in the setting of aSDoH, independent of prevalent obesity.

Despite the large body of literature supporting obesity-related NK dysfunction, the mechanisms by which obesity modulates NK biology have remained unclear. Lipids are thought to play a role; NK cells of individuals with obesity can undergo metabolic reprogramming due to intracellular lipid enrichment and subsequent mitochondrial dysfunction.^36^ While we observed obesity-related NK cell dysfunction, our results did not suggest mitochondrial pathogenesis. This difference could relate to differences in sample processing. We performed our experiments on primary NK cells, while other studies have expanded NK cells prior to analysis. By investigating primary NK cells, we were able to link obesity-related NK cell dysfunction to serum LDL levels. This was based on the association of serum LDL levels with NK cell dysfunction in subjects with obesity, as well as pathological changes in NK phenotype and function after *in vitro* exposure to LDL.

One of our central findings is that LDL reduces NK cytotoxicity and function through DUSP1. This effect was only seen in primary NK cells and not in the IL-2 dependent NK-92 cell line, consistent with the observation that IL-2-mediated NK activation can prevent LDL-mediated repression of cytotoxicity^72^. Nonetheless, our findings reinforce the role of LDL as a potent NK cell immunomodulator. Moreover, our transcriptomic analysis identifies DUSP1 as a common factor linking obesity- and LDL-mediated regulation of NK cells. DUSP1 is a potent anti-inflammatory negative regulator of MAPK-induced NFκB activation^43^, and lipids are reported to induce inflammation through NF-κB activation. Analysis of public datasets have demonstrated *DUSP1* expression in specific NK cell subtypes, indicating a potential role in NK biology.^73, 74^ Hence, our data suggest that DUSP1 may provide a novel mechanistic link between obesity, LDL, and NK cell dysfunction.

Although DUSP1 has not previously been implicated in obesity- or LDL-related immune suppression, it is strongly linked to prevalent obesity.^75^ *Dusp1*^-/-^ mice are resistant to diet-induced obesity,^76^ whereas DUSP1 is upregulated in the subcutaneous adipose tissue, plasma, and PBMCs of individuals with obesity.^77^ This obesity-induced DUSP1 upregulation is reduced by physical activity.^77^ *DUSP1* single nucleotide polymorphisms are also associated with obesity-related metabolic complications.^78^ DUSP1 is also linked to immune cell function in the context of CVD. Specifically, *DUSP1* was the second highest DE gene in monocytes isolated from atherosclerosis patients compared with healthy volunteers.^79^ However, murine models have yielded contradictory results on the role of DUSP1 in atherogenesis. Transferring myeloid *Dusp1*^-/-^ bone marrow into *Ldlr*^-/-^ mice accelerates atherosclerosis,^80^ while germline and myeloid *Dusp1* deletion is protective in *ApoE*^-/-^ mice^81^. Our findings suggest that these differences may be due partly to differences in LDL receptor expression between murine models, although cell-specific DUSP1 regulation may also play a role^82^. DUSP1 and LDL are also reported to regulate each other. *Dusp1*^-/-^ mice have decreased VLDL/LDL levels, ^83^ while modified and oxidized LDL can induce DUSP1 in endothelial cells ^84^ and monocytes.^85^ Together, these findings emphasize the crosstalk between DUSP1 and LDL in immune and stromal cells. Our study fundamentally advances the field by establishing DUSP1 as a central modulator of overweight/obesity and LDL-mediated NK dysfunction in at-risk subjects chronically exposed to aSDoH.

Another key finding of our study is the link between LDL, lysophagy and NK cell dysfunction through DUSP1. We observed that LDL reduced the abundance of intracellular lysosomes in NK cells, which is likely related to induction of lysophagy. LDL has previously been shown to alter lysosomal function and induce lysosomal damage in endothelial cells and macrophages.^52, 86, 87^ In inflammation-induced cardiomyopathy, DUSP1 also regulates mitophagy ^88^, a process of selective autophagy similar to lysophagy. DUSP1 can also inhibit mast cell degranulation^89^, further supporting a potential role in repressing NK cell degranulation. We extend these findings to NK cells, showing that LDL induces lysosomal damage and lysophagy, at least in part through DUSP1. Through this mechanism, DUSP1 represses NK cell degranulation and cytotoxicity. Lysophagy also provides an important mechanistic link to explain how obesity might modulate NF-κB activation through LDL. Lipid treatment of various cells has been reported to activate NFκB, but mainly requires modified or oxidized LDL. Because LDL becomes oxidized upon accumulation in lysosomes ^52^, lysosomal damage is likely to promote oxLDL accumulation, activate NFκB-dependent inflammation, and induce DUSP1 as a negative feedback regulator.

Our results may also help to explain the mechanisms underlying obesity-related malignancy risks. NK cells utilize protrusions to recognize and form immunological synapses with tumor cells, thereby facilitating antitumor cytotoxicity. Tumors can evade NK-mediated cytotoxicity by reducing protrusion abundance on tumor-infiltrating NK cells.^42^ LDL reduced the abundance of NK cell protrusions, which could promote malignancy in individuals with obesity. While future investigations will need to clarify the mechanisms of LDL-driven protrusion loss, our transcriptomic results implicate Rho-GTPases as a potential link. Rho-GTPases mediate cytoskeletal reorganization^90^ and are dysregulated by LDL in NK cells. This suggests that Rho-GTPases might regulate LDL-related NK protrusion loss.

While we have shed light on a unique pathway in NK cell dysregulation in obesity and hyperlipidemia for a population disproportionately impacted by aSDoH, we do acknowledge study limitations. The cross-sectional analyses of human data in our study limits our knowledge on the directionality of the observed associations. Additionally, we were unable to determine if DUSP1 might indeed be regulated by the exposure to chronic stress as aSDoH and if this precedes obesity development. Larger, diverse longitudinal studies are needed at different geographical sites to determine causality. While the inclusion of only AA women could be seen as a limitation, we believe focusing on a group at highest risk for obesity and related cardiometabolic disorders is foundational to identifying targets for future interventions for a population traditionally underrepresented in biomedical research but most impacted by aSDoH.

In summary, our data implicate DUSP1 as a crucial obesity-induced factor that may promote CVD and cancer risk through its immunosuppressive effects. To our knowledge, our study is the first demonstrating a mechanistic link between obesity, LDL, and NK cell function via DUSP1. The study cohort represents a population at highest risk for obesity but historically underrepresented in biomedical research, increasing the potential impact of our work. Future studies should further examine the expression of DUSP1 in an immune subset-specific manner and will need to clarify the role of chronic environmental and psychosocial stressors in obesity-related NK modulation. DUSP1 may also play a role in these processes, as the *DUSP1* gene harbors binding sites for important transcriptional regulators of stress-related immune dysregulation, like NFκB and glucocorticoid receptor.^91^ In the future, DUSP1 may represent a therapeutic target or biomarker in at-risk populations. Future work may involve longitudinal studies of DUSP1 levels in diverse populations most impacted by CVD and cancer disparities. Additionally, community-informed interventions targeting aSDoH, obesity, and hyperlipidemia might impact DUSP1-driven NK cell dysfunction to reduce CVD or cancer risk by promoting physical activity and other lifestyle changes.^77^ Ultimately, this study emphasizes the need for intentional representation of diverse populations in fundamental mechanistic research, and the potential insights gleaned in this approach.

## Acknowledgments

The authors like to thank all the support received by NHLBI, NIMHD, and NIH. In particular, we are grateful for the participants of our clinical trials, our community advisory board, the clinical team supporting our research, and the NIH Bloodbank (especially Dr. West and Dr. Lewis). Special thanks is given to Dr. Dagur and Dr. McCoy at the NHLBI Flow Cytometry and his staff namely Dr. Ankit Saxena, and Dr. Li from the NHLBI DNA Sequencing and Genomics Core and his staff Dr. Liu and Dr. Luo, and Dr. Gucek and his staff from the Proteomics Core as well as the NHLBI Bioinformatics and Computational Biology Core under Dr. Pirooznia’s supervision and the NHLBI Electron Microscopy Core under Dr. Bleck’s supervision. We’d like to thank all colleagues in Dr. Michael Sack’s and Dr. Richard Childs’s laboratories for their outstanding support throughout this study, namely Mr. Shahin Hassanzadeh, Dr. Kim Han, Dr. Joseph Clara, Dr. Kate Stringaris, and Dr. Emily Levy. Furthermore, we’d like to thank Ms. Quan Yu for her technical support in this study.

## Author Contribution

YB and TPW conceptualized the study and wrote the initial manuscript. KS and MP analyzed RNAsequencing and proteomics datasets. KS and DMS analyzed publicly available dataset. YB, ASB, CAG-H, LOW, BST, and YJ performed and analyzed experiments. PD assisted with all flow cytometry-based assays and analysis. LC and MI created the DUSP1 overexpressing and control NK92 cell lines. VMM, BSC, and TPW recruited and worked with the community and patients. DSJA, RNR, and RWC provided support and protocols throughout the study. CKEB and PD provided crucial support for electron microscopy experiments and all flow cytometry. All authors provided critical feedback and edits to the manuscript.

## Declaration of Interest

None.

## Funding

The project was funded by support from the Intramural Research Program of the National Institutes of Health HL006168 and HL006225 to Tiffany M. Powell-Wiley. This research was made possible through the NIH Medical Research Scholars Program, a public-private partnership supported jointly by the NIH and generous contributions to the Foundation for the NIH from the Doris Duke Charitable Foundation, Genentech, the American Association for Dental Research, the Colgate-Palmolive Company, Elsevier, alumni of student research programs, and other individual supporters via contributions to the Foundation for the National Institutes of Health.

## STAR Methods

### Lead Contact

The lead contact for this manuscript is the corresponding author Dr. Tiffany M. Powell-Wiley.

### Materials availability

Upon reasonable request materials can be obtained from the corresponding author. Potentially a MTA has to be established before material transfer can be performed.

### Data and code availability

Upon reasonable request data can be obtained from the corresponding author.

## EXPERIMENTAL MODEL AND SUBJECT DETAILS

### Human Subjects

Study approval was obtained from the Institutional Review Board (IRB) at National Heart, Lung and Blood Institute (NHLBI), National Institutes of Health (NIH) in accordance with the principles of Declaration of Helsinki. The guidelines for good clinical practice and in the Belmont Report (National Commission for the Protection of Human Subjects of Biomedical and Behavioral Research) were followed exactly. Participants were enrolled under the IRB approved clinical trials NCT01143454 and NCT00001846. All study participants provided written informed consent and blood bank donors were de-identified prior to receiving.

#### Cohort 1

Participants (N=29) were AA women from Washington, D.C. metropolitan area. Participants had either no overweight/obesity (NO/O n=15) or had overweight/obesity (O/O n=14) without prevalent cardiovascular disease. Overweight was defined as a body mass index (BMI) ≥25 kg/m^2^ and obesity was defined as a BMI ≥30 kg/m^2^. Individuals were recruited to the Clinical Center at the National Institutes of Health (NIH) and underwent a physical examination, medical history review, and provided blood samples. Whole blood samples underwent flow cytometry-based analysis of blood cells. Additionally, NK cells were isolated, and the NK cell degranulation assay performed as described below (NO/O n=15, O/O: n=13 due to one participant with insufficient blood draw). Patient demographics for cohort 1 are described in **Supplementary Table 1**.

#### Cohort 2

Participants (N=60) were AA adults from the Washington, D.C. metropolitan area. Participants underwent a physical exam, clinical labs, and serum was biobanked. Participant demographics for cohort 2 are provided in **Supplementary Table 2**.

## METHOD DETAILS

### Characterization of whole blood immune cells and NK cell phenotype

Heparinized whole blood (0.5ml) from the study participants was prepared as described previously ^60^. Subsequently two antibody panels were stained and analyzed using flow cytometry. Firstly, a panel generally characterizing immune cell distribution and immune cell-platelet aggregates has been utilized as described previously ^60^. This panel included the following antibody mix, anti-CD3-PECy7 (UCHT1 clone), anti-CD14-APC (61D3 clone), anti-CD15-BV786 (HI98 clone), anti-CD16-BUV395 (3G8 clone), anti-CD19-BV650 (SJ25C1 clone), anti-CD42b-BV421 (HIP1 clone), anti-CD45-PerCP/Cy5.5 (2D1 clone), anti-CD56-FITC (NCAM16.2 clone), anti-CD193-APC/Cy7 (5E8 clone), and anti-CD203c-PE (NP4D6 clone). Afterwards cells were thoroughly washed twice using flow buffer and fixed in flow buffer containing 1% PFA. All samples were analyzed using the LSR Fortessa (BD Bioscience, USA). Results were obtained using FlowJo10 software (FlowJo LLC, USA) with the gating scheme described previously^60^.

### Natural Killer (NK) cell isolation and treatment

PBMCs from peripheral blood from study participants or buffy coats from blood bank donors were isolated using SepMate^TM^ Tubes and Lymphoprep^TM^ separation media (both StemCell Technologies, USA) utilizing the density gradient technique per manufacturers’ recommendation. Next, CD3+ cells (T cells and NKT cells) were removed via positive selection. According to manufacturers’ protocol, freshly isolated PBMCs were tagged with magnetic CD3 MicroBeads (Miltenyi, USA) and then magnetically separated utilizing LS Columns (Miltenyi, USA). kit (Miltenyi, USA) In a similar manner, the flow through (CD3-PBMC) was then subject to a second positive selection step utilizing magnetic CD56 MicroBeads (Miltenyi, USA) to obtain CD3^-^/CD56^+^ NK cells. Isolated NK cells were used after a 30 min resting period in Aim V media (Gibco, USA). Afterwards cells were counted and prepared for treatment immediately.

### Treatments and substances

Isolated NK cells were treated overnight (18h) at 37°C/5% CO_2_ in AimV media (Gibco, USA) with or without the addition of 500μg/ml (50mg/dl) low-density lipoprotein (LDL) (Millipore EMD, USA). The Dusp1 inhibitor (Millipore Sigma, USA, CAS 15982-84-0) was used at 5μM. The MG132 inhibitor (Millipore Sigma, USA) was used at 500nM. Chloroquine (Enzo Life Sciences) was used at 1μM.

### Degranulation assay – NK cell activity

The potential of naïve freshly isolated NK cells to degranulate towards K562 cells was determined using flow cytometry. K562 cells (ATCC) were cultured in RPMI 1640 supplemented with 10% FBS. Prior setup experiments have revealed a 1:1 ratio of freshly isolated naïve NK cell: K562 cells to be optimal for our experimental conditions. 2.5 x 10^5^ NK cells were treated as described above and in each corresponding Figure. After 18h of incubation the treatment was removed and K562 cells were added for a total of 5h to allow NK cell reaction. After 1h of treatment GolgiStop (BD Biosciences, USA) was added per manufacturers’ recommendation to the co-culture to inhibit further cytokine secretion and allow maximum detectable intracellular cytokine labeling for flow cytometry. Afterwards, cells were centrifuged at 300 x g for 5min, the supernatant collected for subsequent ELISA cytokine quantification, and the extracellular proteins CD3, CD56 and CD107a stained using anti-CD3-APC/Cy7 (SK7 clone, BioLegend, USA), anti-CD56-BV450 (NCAM16.2 clone, BD Biosciences, USA), and anti-CD107a-PE (H4A3 clone, BioLegend, USA) stained in flow buffer (1L: PBS pH7.4 with 500μl 0.5M EDTA pH8.0 and 0.2% BSA) for 20min at 4°C in the dark. Then cells were washed twice sing flow buffer, fixed and permeabilized using BD Cytofix/Cytoperm solution as recommended by the manufacturer (BD Bioscience, USA). Afterwards, intracellular cytokines were stained using anti-TNFα-FITC (Mab11 clone, BioLegend), anti-IFNγ-APC (4S.B3, BioLegend, USA), anti-Perforin-PE/Cy7 (dG9 clone, BioLegend), anti-GranzymeB-eFluor450 (N4TL33 clone, Invitrogen, USA), and anti-GM-CSF-PerCP/Cy5.5 (BVD2-21C11 clone BioLegend) for 30min at room temperature in the dark. Cells were then washed and resuspended in flow buffer containing 1% FA to be analyzed using the LSR Fortessa (BD Bioscience, USA). Data were analyzed using FlowJo 10 software (FlowJo LLC, USA). The gating strategy was set by utilizing FMO controls in Cell Activator Cocktail (R&D, USA) treated NK cells.

### *Ex vivo* experimental setup to determine the impact of patient serum on NK cell function

NK cells were isolated as described above from a healthy blood bank donor buffy coat. 2×10^5^ NK cells were treated over night with 10% patient serum (derived and biobanked from Red Top serum tubes without anti-coagulant at their Clinical Center visit) in Aim V media for 18h. Afterwards the NK cell degranulation assay was performed as described above. Afterwards the obtained results per study participant were utilized in statistical models to determine potential significant correlations with clinical markers detected at the participants’ visit date. Due to the nature of this experimental setup all results are obtained in a blinded fashion.

### Cytokine ELISA

The two cytokines IFNγ and TNFα were detected from cell culture supernatants of the degranulation assays utilizing the respective R&D DuoSet (R&D Systems, USA) ELISAs per manufacturers’ recommendation.

### Scanning Electron Microscopy (SEM) and Transmission Electron Microscopy (TEM)

NK cells were treated as indicated and immediately fixed in 2.5% glutaraldehyde (0.1M calcium chloride, 0.1M sodium cacodylate buffer, pH 7.2). Samples were prepared on 0.1μm filters as described previously.^60, 92, 93^ Gold/palladium sputter coated NK cells were imaged using the Hitachi S-3400N1 SEM at 7.5keV and the JOEL JEM1400 transmission electron microscope (NHLBI Electron Microscopy Core).

### RNA Sequencing and bioinformatics analysis of NK cells from participants with or without overweight/obesity as well as control or LDL exposed NK cells

RNA sequencing was performed from freshly isolated primary NK cells isolated as described above from participants’ peripheral blood (n=5 each lean or obese). Healthy donor buffy coats treated with vehicle (control) or LDL (n=4 sets). After isolation and/or treatment, NK cells were lysed in Trizol (15596026, Thermo Fisher Scientific, USA). RNA extraction was performed using the RNeasy Mini Kit (74104, Qiagen, USA) per manufacturers’ recommendation. An additional step of DNase I treatment was performed, and cells were washed twice thereafter. Concentration was determined using NanoDrop Spectrophotometer. Libraries were prepared usingL---Lkit (Illumina) and were sequenced on Illumina Novaseq for paired-end 100bps using standard Illumina sequencing primers. RNA seq fasta files quality was checked using theLFastQC (http://www.bioinformatics.babraham.ac.uk/projects/fastqc). The adapters were trimmed using Trimmomatic.^82^ The RNA sequence data was aligned to the human genome (GRCh38) usingLSplice aware Transcripts Aligner (STAR).^83^LfeatureCounts^84^ was used for gene expression quantification andLLimma-voom^85^ was used to perform differential expression analysis. Genes with p values <0.05 were considered differentially expressed (DE). Pathway enrichment analysis was performed on DE genes using clusterProfiler.^86^ Pathways with q values <0.05 were considered significant enrichment except for pathway results from concordant DE genes, where p value significant pathways were reported.LAdditionally, a custom gene matrix transposed (gmt) file comprising of 71 pathways were generated and fisher test was performed on DE gene list in R and the false discovery rate (FDR) corrected p value is reported.LR (R Core Team (2017). R: A language and environment for statistical computing. RL(Foundation for Statistical Computing, Vienna, Austria. URLLhttps://www.R-project.org/) statistical software was used for data visualization to perform principal component analysis (PCA).

### Proteomics

Protein extraction and digestion: Proteins from each group of samples are being quantified using the BCA assay Kit. Twenty microgram of protein was denatured in 8M Urea,25mM TEAB buffer followed by reduction (DTT) and alkylation (iodoacetamide) and then the alkylated proteins were diluted to <1M Urea buffer using 25mM TEAB, pH 8 buffer before performing Trypsin digestion overnight. Each sample was digested overnight using Trypsin enzyme (enzyme to substrate ratio of 1:12.5) to get the peptide solution. Trypsin will cleave proteins at lysine and arginine amino acid residues yielding tryptic peptides. The samples were acidified the next day using formic acid to bring pH<3 to stop the trypsin activity and were processed through Zip-Tip using C18 tips (Millipore) to clean-up and concentrate the peptides before going through mass spec analysis.

Liquid Chromatography – Tandem MS Analysis: Protein identification by LC-MS/MS analysis of peptides was performed using an Orbitrap Fusion Lumos Tribid mass spectrometer (Thermo Fisher Scientific, San Jose, CA) interfaced with an Ultimate 3000 Nano-HPLC apparatus (Thermo Fisher Scientific, San Jose, CA). Peptides were fractionated by EASY-Spray PepMAP RPLC C18 column (2m, 100A, 75 m x 50cm) using a 120-min linear gradient of 5-35% ACN in 0.1% FA at a flow rate of 300 nl/min. The instrument was operated in data-dependent acquisition mode (DDA) using FT mass analyzer for one survey MS scan on selecting precursor ions followed by 3 second data-dependent HCD-MS/MS scans for precursor peptides with 2-7 charged ions above a threshold ion count of 10,000 with normalized collision energy of 37%. Survey scans of peptide precursors from 300 to 2000 m/z were performed at 120k resolution and MS/MS scans were acquired at 50,000 resolution with a mass range m/z 100-2000.

Protein Identification and data analysis: All MS and MS/MS raw spectra from each set were processed and searched using Sequest HT algorithm within the Proteome Discoverer 2.2 (PD2.2 software, Thermo Scientific).The settings for precursor mass tolerance was set at 12 ppm, fragment ion mass tolerance to 0.05 Da, trypsin enzyme with 2 mis cleavages with carbamidomethylation of cysteine as fixed modifications, deamidation of glutamine and asparagine, oxidation of methionine as variable modifications. The Human sequence database from Swiss sprot was used for database search. Identified peptides were filtered for maximum 1% FDR using the Percolator algorithm in PD 2.2 along with additional peptide confidence set to medium. The final lists of protein identification/quantitation were filtered by PD 2.2 with at least 2 unique peptides per protein identified with medium confidence. For the quantitation, label-free approach has been used, where the area under the curve for the precursor ions is used to calculate the relative fold change between different peptide ions.

### Flow Cytometry analysis

NK cells were treated as indicated, centrifuged at 300 x g for 5min, and the supernatant was discarded. Next, NK cells were washed twice using flow buffer, fixed and permeabilized using BD Cytofix/Cytoperm solution as recommended by the manufacturer (BD Bioscience, USA). Subsequently, NK cells were incubated with the according antibody cocktail for 30min at RT in the dark.

To determine Dusp1 expression levels the following antibody panel was used: anti-Lamp1-AF488 (clone H4A3, BioLegend, USA) and anti-Dusp1-AF594 (clone E6)

To determine galectin-3 or galectin-8 expression the following: intracellular protein and lysosomes were stained using anti-Lamp1-BV421 (clone H4A3, BioLegend, USA), anti-Galectin-3-AF488 (clone B2C10, Santa Cruz Biotechnologies, USA), and galectin-8-AF790 (clone C-8, Santa Cruz Biotechnologies, USA)

After antibody incubation cells were washed and resuspended in flow buffer with 1% PFA. Flow cytometry was performed using the LSR Fortessa (BD Bioscience, USA) and data were analyzed using FlowJo 10 software (FlowJo LLC, USA).

### Luminex assay for MAPK and phosphorylation

NK cells were treated as indicated, centrifuged at 300 x g for 5min, and the supernatant was discarded. Subsequently cells were lysed using MilliplexMAP Cell Signaling Universal Lysis Buffer. Luminex technology was used to quantify presence of phosphorylated MAPK signaling proteins, their unphosphorylated counterparts, and tubulin as housekeeping protein. The assay was conducted as per the manufacturers’ recommendation.

### Immunofluorescence

Cells were incubated as indicated, centrifuged at 300 x g for 5min, and the supernatant was discarded. Subsequently cells were fixed and permeabilized using the BD Cytofix/Cytoperm solution as recommended by the manufacturer (BD Bioscience, USA). Afterwards cells were washed and incubated with the antibody staining cocktails for 1hour at RT in the dark.

F-actin was labeled using Phalloidin-AF488 (ThermoFisher, USA) while lysosomes were labeled using anti-Lamp1-AF594 (clone H4A3, Santa Cruz Biotechnologies, USA). Additionally, nuclei were labled using DAPI solution.

### NK92 Dusp1 overexpressing cell line creation

Dusp1 was expressed in NK92 cells by the Sleeping Beauty transposon system. An empty vector control cell line was created as corresponding control.

Sleeping Beauty vectors construction: The backbone transposon plasmid pT2/HB was a gift from Perry Hackett (Addgene plasmid # 26557). The transposase plasmid pCMV(CAT)T7-SB100 was a gift from Zsuzsanna Izsvak (Addgene plasmid # 34879). The DUSP1 gene ORF (Ensembl, ENSG00000120129) was synthesized from GeneArt (Thermo Fisher) and was cloned into pT2/HB vector through In-Fusion HD Cloning Plus kit (Takara Bio) following manufacturer’s instructions.

Electroporation of NK-92 cells: NK-92 cells were harvested, counted, and resuspended in 100 uL Cell Line Nucleofector Solution R (Lonza) at the concentration of 1×10^8^ cells/mL. pT2/HB transposon plasmids encoding DUSP1 or truncated CD34 (16 ug) and pCMV(CAT)T7-SB100 transposase plasmids (4ug) were added for each electroporation based on optimization experiments. The electroporation was performed on Nucleofector 2b (Lonza) using the program A-024. Cells were returned to incubator 48 hours and resuspended at desired concentration with fresh complete medium. On day 7 post electroporation, CD34+ cells were sorted using anti-human CD34 magnetic beads (Miltenyi). The percentage of CD34+ cells before and after sorting was measured using another anti-human CD34 mAb (Miltenyi, clone AC136) binding a different epitope than the one recognized by anti-human CD34 magnetic beads. The sorted cells were expanded and used for phenotyping and functional assays.

## Quantification and Statistical Analysis

### Statistical analysis

All wet lab derived data were analyzed using PRISM 7.0 (GraphPad) and Microsoft Excel software. Each dataset was tested for normality using the Shapiro-wilk Test to determine subsequent statistical models. All significance thresholds were designated at P values < 0.05. Depending on the nature of the dataset, paired analysis was performed. Normally distributed datasets were analyzed using t-test, paired t-test, One-way ANOVA with Dunnett correction for multiple comparison (no pairing), or Mixed-effects analysis with Geisser-Greenhouse correction and Dunn’s correction for multiple comparison analysis (paired). Nonparametric datasets were analyzed using Mann-Whitney test, Kruskal-Wallis test (unpaired) or Friedman test (paired) either with Dunn’s correction for multiple comparison. All data is presented as mean ± standard deviation of the mean and the exact sample size (n) is provided in the figure legends.

Multivariable regression analysis was performed using STATA release 12 (StataCorp., College Station, Tx, USA). Unadjusted and adjusted multivariable linear regression modeling were used to evaluate the relationship between psychosocial measures, clinical parameters, and ex vivo characteristics. P values of <0.05 are reported as statistically significant. All these analyses were performed in a blinded fashion due to the nature of the experiment.

## KEY RESOURCES TABLE

| REAGENT or RESOURCE | SOURCE | IDENTIFIER |
| --- | --- | --- |
| **Antibodies** |  |  |
| Anti-CD3-PECy7 | EBioscience | RRID: AB_1582253 |
| Anti-CD14-APC | EBioscience | RRID: AB_10669167 |
| Anti-CD15-BV786 | BD Biosciences | RRID: AB_2738444 |
| Anti-CD16-BUV395 | BD Biosciences | RRID: AB_2744293 |
| Anti-CD19-BV650 | BD Biosciences | RRID: AB_2744313 |
| Anti-CD42b-BV421 | BioLegend | RRID: AB_2715931 |
| Anti-CD45-PerCP/Cy5.5 | BD Biosciences | RRID: AB_400194 |
| Anti-CD56-FITC | BD Biosciences | RRID: AB_400025 |
| Anti-CD193- APC/Cy7 | BioLegend | RRID: AB_10898162 |
| Anti-CD203c-PE | BioLegend | RRID: AB_756044 |
| Anti-CD3- APC/Cy7 | BioLegend | RRID: AB_10645474 |
| Anti-CD56-BV650 | BD Bioscience | RRID: AB_2738568 |
| Anti-CD107a-PE | BioLegend | RRID: AB_1186040 |
| Anti-TNFα-FITC | BioLegend | RRID: AB_315258 |
| Anti-IFNγ-APC | BioLegend | RRID: AB_315237 |
| Anti-Perforin-PE/Cy7 | BioLegend | RRID: AB_2572048 |
| Anti-GranzymeB-eFluor450 | Invitrogen | RRID: AB_2724392 |
| Anti-GM-CSF-PerCP/Cy5.5 | BioLegend | RRID: AB_11146993 |
| Anti-Galectin-3-AF488 | Santa Cruz<br>Biotechnologies | Catalog # sc-32790 AF594 |
| Anti-Galectin-8-AF790 |  | Catalog # sc-377133 AF790 |
| Anti-Lamp1-BV421 | BioLegend | Catalog # 328626 |
| Anti-Lamp1-AF488 | BioLegend | Catalog # 328610 |
| Anti-Dusp1-AF594 | Santa Cruz<br>Biotechnologies | Catalog # sc-373841 AF594 |
| Phalloidin-AF488 | ThermoFisher | Catalog # A12379 |
| Anti-Lamp1-AF594 | Santa Cruz<br>Biotechnologies | Catalog # sc- AF594 |
| Anti-CD34-PE | BioLegend | Catalog # 343505 |
| **Bacterial and virus strains** |  |  |
| Stellar Competent Cells | Takara Bio | Catalog # 636766 |
| **Chemicals, peptides, and recombinant proteins** |  |  |
| MACSQuant® Running Buffer | Miltenyi | Catalog # 130-092-7470 |
| CD3 MicroBeads, human | Miltenyi | Catalog # 130-050-101 |
| CD56 MicroBeads, human | Miltenyi | Catalog # 130-050-401 |
| 20% Paraformaldehyde aqueous solution | Electron<br>Microscopy<br>Sciences | Catalog # 50-980-493 |
| SepMate™-50 (IVD) | StemCell<br>Technologies | Catalog # 85450 |
| Lymphoprep<sup>TM</sup> | StemCell Technologies | Catalog # 07851 |
| Aim V media | Gibco | Catalog # 12055091 |
| Low-density lipoprotein (LDL) | Kalen Biomedical | Catalog # 770200-8 |
| Dusp1 inhibitor | Millipore Sigma | Catalog # 317496-10MG |
| MG132 | Millipore Sigma | Catalog # 474791-1MG |
| Chloroquine | Tocris | Catalog # 4109 |
| RPMI 1640 | Gibco | Catalog # 11875093 |
| FBS, heat inactivated | VWR | Catalog # 97068-091 |
| Horse Serum | Millipore Sigma | Catalog # H1270-500 |
| IL-2 (Tecin<sup>TM</sup>) | National Cancer Institute | Ro 23-6019 |
| GlutaMax | Gibco | Catalog # 35050061 |
| GolgiStop | BD Biosciences | Catalog # 554724 |
| BD Cytofix/Cytoperm | BD Biosciences | Catalog # 554722 |
| Cell Activator Cocktail | Tocris | Catalog # 5476 |
| IFNγ R&D DuoSet ELISA | R&D Systems, USA | Catalog # DY285B-05 |
| TNFα R&D DuoSet ELISA | R&D Systems, USA | Catalog # DY210-05 |
| Trizol | Thermo Fisher Scientific | Catalog # 15596026 |
| RNase-Free DNase Set | Qiagen | Catalog # 79254 |
| RNeasy Mini Kit | Qiagen | Catalog # 74104 |
| MilliplexMAP Cell Signaling Universal Lysis Buffer | Millipore Sigma | Catalog # 43-040 |
| **Critical commercial assays** |  |  |
| 9plx Multipathway Total Kit | Millipore Sigma | Catalog # 48-681MAG |
| MilliplexMAP 9-Plex Multi-Pathway Magnetic Bead Signaling Kit | Millipore Sigma | Catalog # 48-680MAG |
| Total b-Tubulin Magnetic Bead MAPmate | Millipore Sigma | Catalog # 46-713MAG |
| **Deposited data** |  |  |
| RNA sequencing NO/O versus O/O | Gene Expression Omnibus | Data will be available with restrictions on Geo and upon reasonable request to the corresponding author. |
| RNA sequencing ctr versus LDL |  |  |
| RNA sequencing Dusp1oe NK92 cells |  |  |
| Proteomics ctr versus LDL |  |  |
| Proteomics Dusp1oe NK92 cells |  |  |
| **Experimental models: Cell lines** |  |  |
| K562 cells (generous gift by Dr. Childs laboratory, NHLBI) |  |  |
| NK92 cells (generous gift by Dr. Childs laboratory, NHLBI) |  |  |
| **Recombinant DNA** |  |  |
| pT2/HB | Addgene | Catalog # 26557 |
| pCMV(CAT)T7-SB100 | Addgene | Catalog # 34879 |
| **Software and algorithms** |  |  |
| FlowJo 10 | FlowJo LLC |  |
| ImageJ | ImageJ |  |
| PRISM 7.0 (GraphPad | GraphPad |  |
| STATA release 12 (StataCorp) | StataCorp |  |

## Notes

### Competing Interest Statement

The authors have declared no competing interest.

