## Supplemental Data and Information for "Social Determinants modulate NK cell activity via obesity, LDL, and DUSP1 signaling"

**Supplementary Tables and Figures:**

**Supplementary Table 1: Patient demographics of Cohort 1 utilized for NK peripheral blood characterization studies.**

| | **All Participants (n=29)** | **Subgroup with PSES measures available (n=17)** |
|---|---|---|
| Sex, Female | 29 (100) | 17 (100) |
| Ethnicity, African American | 29 (100) | 17 (100) |
| Age, years | 54.79 ± 15.8 | 54.02 ± 13.6 |
| Body Mass Index (BMI), $kg/m^2$ | 32.11 ± 10.0 | 33.96 ± 8.9 |
| ASCVD, 10-yr risk score[†] | 7.70 ± 6.1 [^] | 7.23 ± 5.9 |
| **Medical History:** | | |
| Hypertension | 10 (34.4) | 7 (41.2) |
| Diabetes | 4 (13.8) | 2 (11.8) |
| Hyperlipidemia on Lipid-lowering Rx | 6 (20.7) | 5 (29.4) |
| **Laboratory Markers:** | | |
| Total Cholesterol (TC), mg/dl | 165.00 ± 31.7 | 181.47 ± 25.7 |
| LDL, mg/dl | 85.41 ± 27.8 | 100.18 ± 24.6 |
| ApoB, mg/dl | 67.26 ± 24.7 | 87.67 ± 18.2 |
| Triglycerides (TG), mg/dl | 63.34 ± 25.9 | 75.94 ± 21.0 |
| HDL, mg/dl | 66.89 ± 20.2 | 66.00 ± 17.6 |
| ApoA1, mg/dl | 150.64 ± 31.2 | 166.33 ± 26.7 |
| **Psychosocial and environmental stressors:** | | |
| NDI[^] | --- | -1.11 ± 2.9 |
| Perceived Neighborhood Physical/Social Environment[&] | --- | 22.00 ± 6.2 |
| Perceived Neighborhood Social Cohesion[#] | --- | 14.00 ± 3.1 |
| Categorical SES, low income[‡] | --- | 5 (29.4) |
| Continuous SES, income /$10k[‡] | --- | 67.65 ± 30.5 |
| Social Isolation[‖] | --- | 0.36 ± 0.7 |
| Depressive Symptoms and Symptom sub-types: [§] | | |
| Overall Depressive Symptoms | --- | 6.53 ± 6.8 |
| Sadness | --- | 1.53 ± 2.0 |
| Anhedonia/Loss of Interest | --- | 0.53 ± 1.2 |
| Lack of Appetite | --- | 0.13 ± 0.4 |
| Lack of Sleep | --- | 1.87 ± 1.8 |
| Change in Thinking | --- | 0.75 ± 1.0 |
| Guilt | --- | 0.40 ±1.1 |
| Tired/Fatigue | --- | 0.93 ±1.2 |
| Movement/Agitation | --- | 0.33 ± 0.9 |

Categorical variables expressed as number of participants followed by the percent of the total (%). Continuous variables expressed as mean ± standard deviation (SD).

[†]ASCVD 10-year risk score – Atherosclerosis Cardiovascular Disease 10-year risk score

[^] NDI - neighborhood deprivation index derived from U.S. Census data (higher score=more socioeconomically deprived neighborhood)

[&] Perceptions about neighborhood physical/social environment measured by validated survey (higher score=greater perceived resources in the physical/social environment)

[#] Perceived neighborhood social cohesion measured by validated survey (higher score=more neighborhood cohesion)

[‡] SES – individual-level socioeconomic status measured (higher value=higher self-reported household income); income<$60,000 = low SES, while income>60,000 = high SES per median household income in the Washington, D.C. area

[||] Social Isolation - measured by validated survey (higher score=increasing social isolation)

[§] Depression and depression sub-scores were measured by validated surveys (higher score=more depression, sadness, lack of appetite, lack of sleep, increased perception of guilt)

**Supplementary Table 2: Patient demographics for Cohort 2 used for *ex vivo* serum component studies.**

| | **Participants (N=60)** |
|---|---|
| **Sex, Female** | 56 (93) |
| **Ethnicity, African American** | 60 (100) |
| **Age, years** | 60.83 ± 10.5 |
| **BMI, kg/m$^2$** | 33.00 ± 7.9 |
| **ASCVD, 10-yr risk score** | 10.75 ± 8.5 |
| **Hypertension** | 38 (63.3) |
| **Diabetes** | 13 (21.7) |
| **Lipid-lowering therapy** | 22 (37) |
| **TC, mg/dl** | 188.98 ± 35.20 |
| **HDL, mg/dl** | 66.57 ± 20.58 |
| **LDL, mg/dl** | 105.5 ± 33.03 |
| **ApoB, mg/dl** | 92.28 ± 24.21 |
| **TG, mg/dl** | 84.97 ± 26.43 |
| **HDL, mg/dl** | 66.57 ± 20.58 |
| **ApoA1, mg/dl** | 169.35 ± 30.29 |

Categorical variables expressed as number of participants followed by the percent of the total (%). Continuous variables expressed as mean ± standard deviation (SD).

**Supplementary Table 3:** In an *ex vivo* approach healthy donor primary NK cells were incubated with donor serum (patients summarized in Supplementary Table 2) for 24hours and afterwards subjected to the degranulation assay. Regression associations between detected CD107a surfaces expression as well as intracellular TNFα and IFNγ expression and clinical parameters in unadjusted, fully adjusted model 1 (adjusted for ASCVD and BMI) and adjusted model 2 (adjusted for ASCVD, BMI, SES) shown as standardized ß-value with the corresponding p-value in parenthesis for all study participants independent of status on lipid-lowering therapies.

| | **CD107a** | | | **TNFα** | | | **IFNγ** | | |
|---|---|---|---|---|---|---|---|---|---|
| | **Unadj.** | **Adj.[1]** | **Adj.[2]** | **Unadj.** | **Adj.[1]** | **Adj.[2]** | **Unadj.** | **Adj.[1]** | **Adj.[2]** |
| **TC** | -0.15 (0.26) | -0.15 (0.25) | -0.20 (0.21) | 0.06 (0.64) | 0.02 (0.87) | 0.07 (0.64) | **-0.37 (0.004)*** | **-0.34 (0.009)*** | **-0.33 (0.03)*** |
| **LDL** | -0.21 (0.11) | -0.22 (0.09) | -0.22 (0.15) | 0.12 (0.35) | 0.08 (0.57) | 0.14 (0.37) | **-0.37 (0.004)*** | **-0.35 (0.008)*** | -0.29 (0.07) |
| **ApoB** | -0.19 (0.17) | -0.21 (0.15) | -0.23 (0.15) | 0.02 (0.89) | -0.01 (0.95) | 0.11 (0.49) | **-0.28 (0.04)*** | *-0.27 (0.05)* | -0.29 (0.08) |
| **TG** | 0.04 (0.73) | 0.04 (0.77) | -0.05 (0.74) | 0.16 (0.21) | 0.17 (0.24) | **0.33 (0.03)*** | -0.04 (0.79) | -0.04 (0.77) | 0.01 (0.93) |
| **HDL** | 0.07 (0.59) | 0.08 (0.53) | 0.06 (0.69) | -0.13 (0.33) | -0.13 (0.34) | -0.22 (0.14) | -0.03 (0.84) | -0.00 (0.95) | -0.11 (0.45) |
| **ApoA1** | 0.05 (0.71) | 0.06 (0.68) | -0.01 (0.93) | -0.09 (0.51) | 0.10 (0.49) | -0.13 (0.43) | -0.06 (0.69) | -0.05 (0.71) | -0.12 (0.47) |

### Supplementary Figures

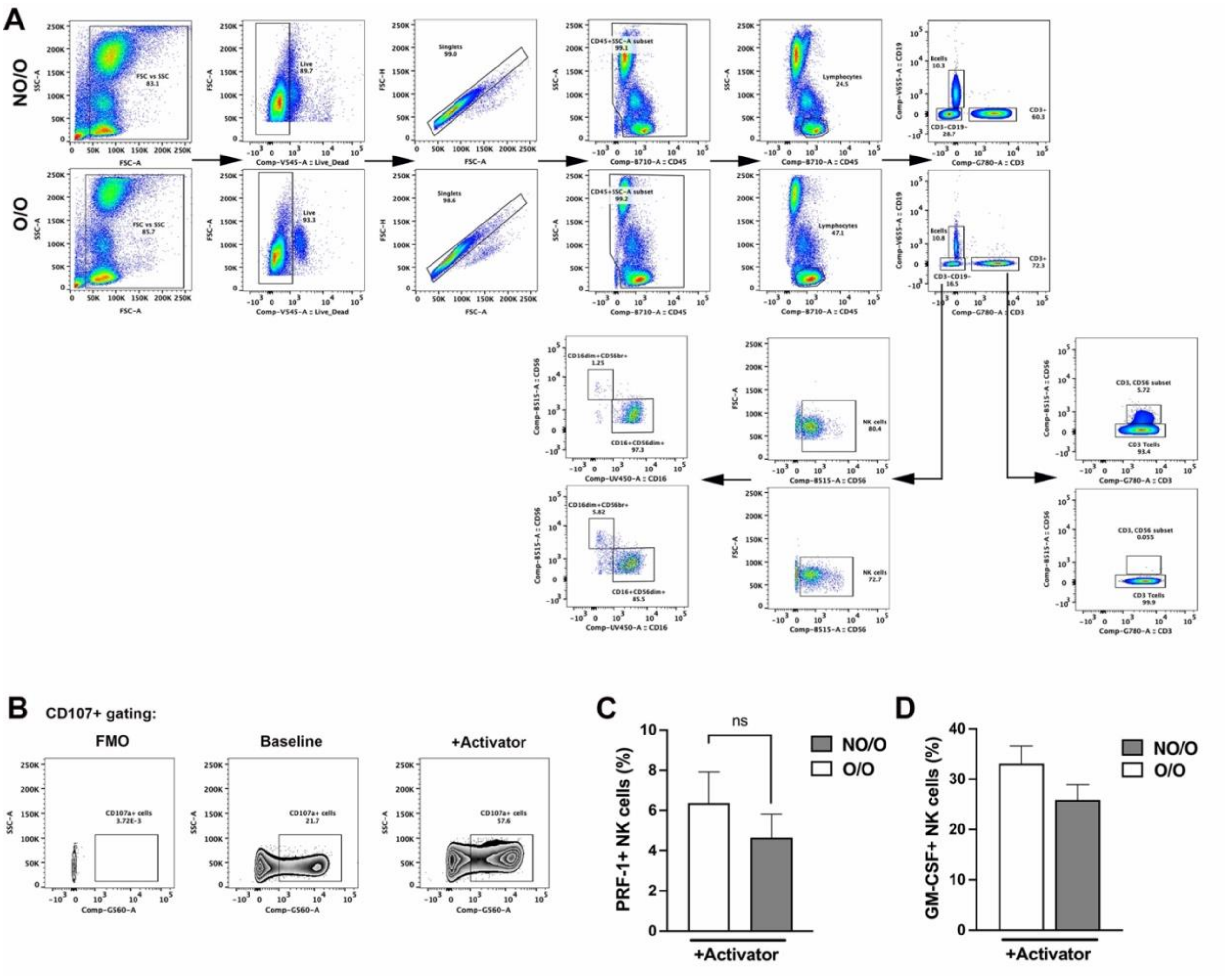


**Supplementary Figure 1: NK cells from individuals with overweight/obesity displayed altered intracellular cytokine expression.**

(**A**) Gating scheme for Figure 1A/B displaying one representative sample of a NO/O and an O/O study participant.

(**B**) Gating scheme with FMO control for CD107a expression.

(**C/D**) Freshly isolated NK cells from NO/O or O/O AA females were subjected to the degranulation assay with subsequent flow cytometry-based analysis of intracellular perforin 1 (**C**), or GM-CSF (**D**) expression. (Significance was established when a p-value $< 0.05$ was present comparing individual groups with the data appropriate test; *indicates significance between groups; A: Mann-Whitney test, B/C: Unpaired T-test)

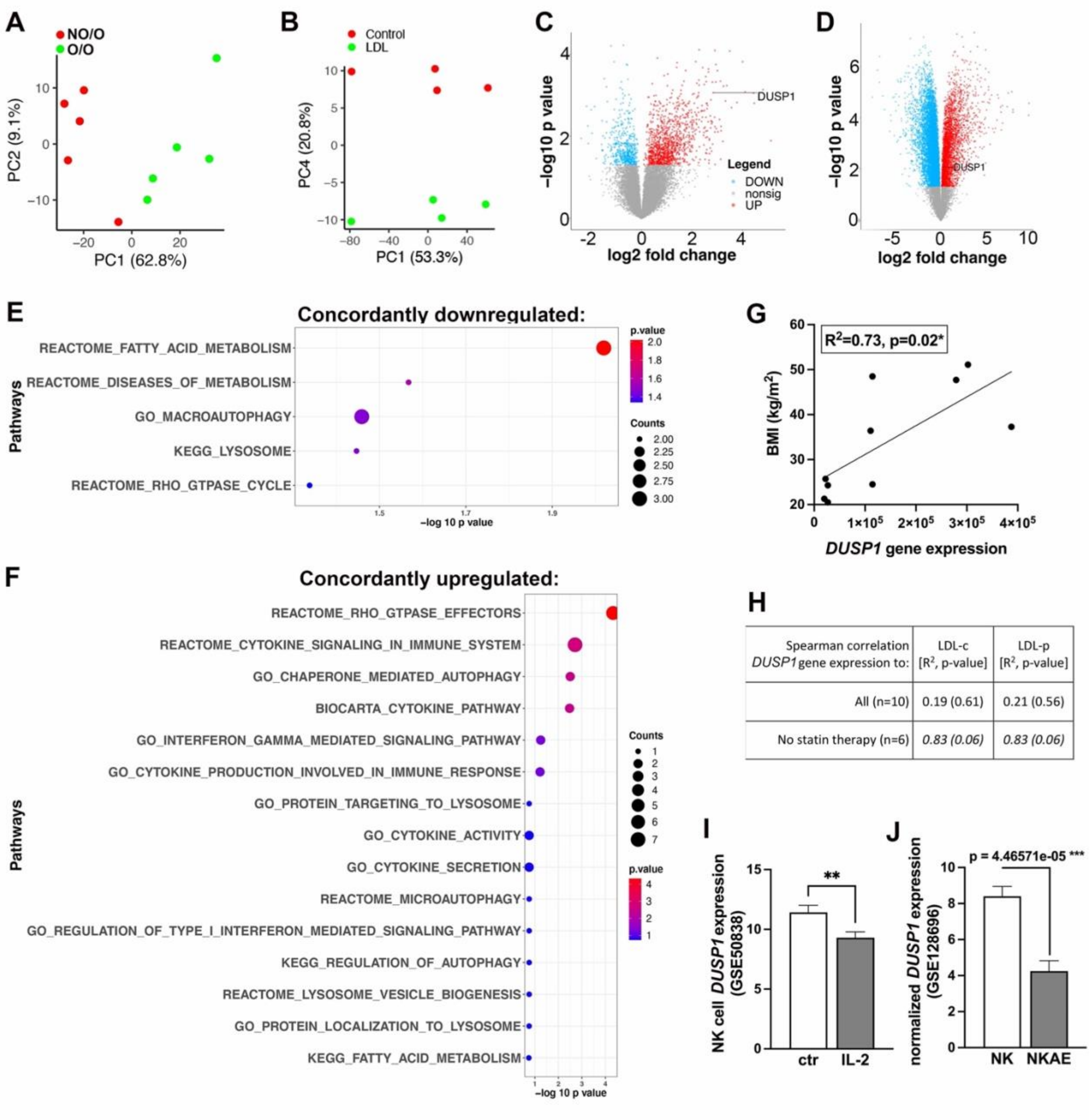


| Spearman correlation *DUSP1* gene expression to: | LDL-c [$R^2$, p-value] | LDL-p [$R^2$, p-value] |
|---|---|---|
| All (n=10) | 0.19 (0.61) | 0.21 (0.56) |
| No statin therapy (n=6) | *0.83 (0.06)* | *0.83 (0.06)* |

**Supplementary Figure 2: Identifying Dusp1 as a commonly regulated gene of potential crucial importance for NK cell function.** Freshly isolated NK cells of NO/O or O/O (n=5 each group) study participants or from healthy donors with subsequent vehicle/LDL exposure (n=4) were subjected to RNA sequencing analysis.

(**A/B**) Principal component analysis (PCA) computed from the differentially expressed genes identified in the NO/O vs O/O (A) or control vs. LDL (B) datasets. Each dot represents a sample from indicated group.

(**C/D**) Volcano plot of genes profiled in N (C) or control vs. LDL (D) highlights the statistically significant (P value <0.05) genes as red (upregulated) and blue (downregulated) in obese or LDL treatment. Remaining genes are shown in grey color. The gene data point representing DUSP1 is highlighted.

(**E/F**) **)** Top pathways enriched by the differentially expressed genes that were concordantly either upregulated (A) or downregulated (B) in in NO/O vs. O/O and control vs. LDL datasets. Each dot is a pathway labeled on y-axis, the size of the dot is scaled to the number of concordant DE genes overlapping with the pathway and are plotted on negative log 10 p value scale on the x-axis.

(**G**) *DUSP1* gene expression levels were correlated with the study participants BMI utilizing Spearman correlation. (n=10)

(**H**) *DUSP1* gene expression levels were subjected to Spearman correlation against LDL-c (concentration) and LDL-p (particle number).

(**I/J**) Two publicly available datasets on IL-2 treated activated NK cells (GSE50838) as well as activated/expanded NK cells (NKAE, GSE128696) were analyzed for potential changes in *DUSP1* expression to provide evidence that *DUSP1* gene expression is affected when NK cell activity is altered. (Significance was established when a p-value < 0.05 was present comparing individual groups with the data appropriate test; *indicates significance between groups; A: unpaired t-test, B/C: Mann-Whitney test)

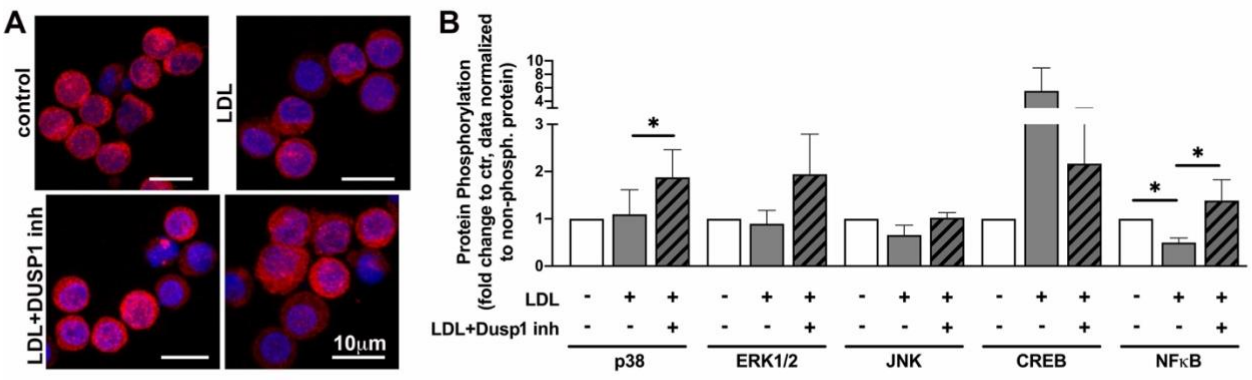


**Supplementary Figure 3: LDL treatment of NK cells regulates NFκB activity in a Dusp1 dependent manner.** Freshly isolated primary NK cells were treated as indicated and subjected to immunofluorescence analysis of NFκB (A, n=3, red=NFκB, blue=nucleus) or a Luminex assay determining MAPK and NFκB pathway activation (B). (Significance was established when a p-value < 0.05 was present comparing individual groups with the Friedman's test and Dunn's correction; *indicates significance between groups; Abbreviations: CREB=cAMP response element, ERK1/2=extracellular signal-regulated protein kinase 1/2, JNK=c-Jun N-terminal kinase, NFκB=Nuclear factor kappa B, p38=p38 mitogen-activated protein kinase)

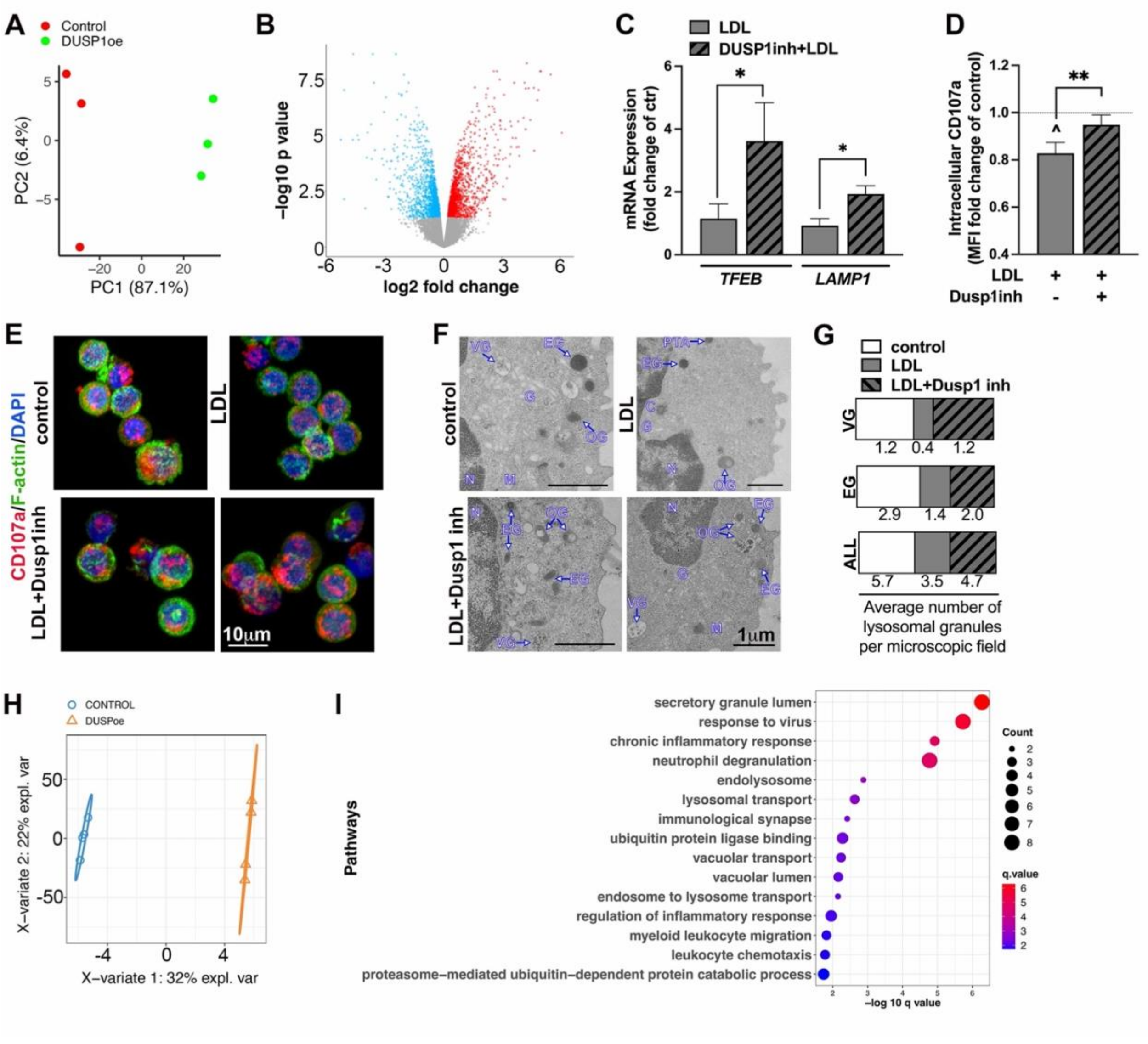

**Supplementary Figure 4: Dusp1 is an important regulator of lysosomal biology.**
(**A**) Principal component analysis (PCA) computed from the differentially expressed genes identified in DUSP1 overexpressing (oe) NK92 cells and their respective empty vector control. Each dot represents a sample from indicated group.
(**B**) Volcano plot of genes profiled in DUSP1 overexpressing (oe) NK92 cells and their respective empty vector control NK92 cells highlights the statistically significant (P value <0.05) genes as red (upregulated) and blue (downregulated) in DUSP1oe NK92 cells.
(**C-G**) Freshly isolated primary NK cells were treated with vehicle, LDL, or LDL in presence of the Dusp1 inhibitor.
(**C**) Freshly isolated primary NK cells were treated with vehicle, LDL, or LDL in presence of the Dusp1 inhibitor. RT-qPCR was performed to determine the impact of LDL and its dependency on Dusp1 on mRNA levels of *TFEB*, a transcription factor for lysosomal biogenesis, and *LAMP1*. (TFEB: n=8, Friedman test with Dunn correction, LAMP1: n=7, RM One-Way ANOVA with Tukey correction)

(**D**) Freshly isolated primary NK cells were treated with vehicle, LDL, or LDL in presence of the Dusp1 inhibitor. Flow cytometry was used to quantify intracellular CD107a expression. (n=10, RM One-Way ANOVA with Sidak correction).
(**E**) Freshly isolated primary NK cells were treated with vehicle, LDL, or LDL in presence of the Dusp1 inhibitor. Immunofluorescence analysis of Lamp-1/CD107a (red) to determine the presence of intracellular lysosomes, F-actin (green), and DAPI to label nuclei (Blue) (n=4).
(**F/G**) Freshly isolated primary NK cells were treated with vehicle, LDL, or LDL in presence of the Dusp1 inhibitor. Transmission electron microscopy analysis of pooled samples from 3 experimental setups. Number of vesicular granules (VG), electron dense granules (EG), as well as all granules (VG, EG, and other granules (OG)) were counted per microscopic field and graphed per treatment condition in (**G**), while representative images are shown in (**F**).
(**H/I**) Proteomics analysis of DUSP1 overexpressing (oe) NK92 cells and their respective empty vector control NK92 cells.
(**H**) Partial least square discriminant analysis (PLSDA) plot of the proteomics data of DUSP1 overexpressing (oe) NK92 cells and their respective empty vector control NK92 cells.
(**I**) Top 15 pathways enriched by differentially expressed proteins in DUSP! Oe NK92 cells vs. empty vector NK92 cells. Each dot is a pathway labeled on y-axis, the size of the dot is scaled to the number of DE genes overlapping with the pathway and are plotted on negative log 10 q value on the x-axis.
(Significance was established when a p-value < 0.05 was present comparing individual groups; *indicates significance between groups; ^indicates significance to vehicle control treatment)
